# Ataxin-3 links NOD2 and TLR2 mediated innate immune sensing and metabolism in myeloid cells

**DOI:** 10.1101/595637

**Authors:** Thomas Chapman, Daniele Corridoni, Seiji Shiraishi, Sumeet Pandey, Anna Aulicino, Simon Wigfield, Maria do Carmo Costa, Marie-Laëtitia Thézénas, Henry Paulson, Roman Fischer, Benedikt M Kessler, Alison Simmons

**Affiliations:** MRC Human Immunology Unit, MRC Weatherall Institute of Molecular Medicine, John Radcliffe Hospital, University of Oxford, Oxford, OX3 9DS, UK; Translational Gastroenterology Unit, John Radcliffe Hospital, University of Oxford, Oxford OX3 9DS, UK; Target Discovery Institute, Nuffield Department of Medicine, University of Oxford, Oxford OX3 7FZ, UK; Department of Neurology, University of Michigan, Ann Arbor, MI, USA; Department of Oncology, University of Oxford, OX3 7DQ, UK

## Abstract

The interplay between NOD2 and TLR2 following recognition of components of the bacterial cell wall peptidoglycan is well established, however their role in redirecting metabolic pathways in myeloid cells to degrade pathogens and mount antigen presentation remains unclear. We show NOD2 and TLR2 mediate phosphorylation of the deubiquitinase ataxin-3 via RIPK2 and TBK1. In myeloid cells ataxin-3 associates with the mitochondrial cristae protein MIC60, and is required for oxidative phosphorylation. Depletion of ataxin-3 leads to impaired induction of mitochondrial reactive oxygen species (mROS) and defective bacterial killing. A mass spectrometry analysis of NOD2/TLR2 triggered ataxin-3 deubiquitination targets revealed immunometabolic regulators, including HIF-1α and LAMTOR1 that may contribute to these effects. Thus, we define how ataxin-3 plays an essential role in NOD2 and TLR2 sensing and effector functions in myeloid cells.

**Significance Statement:** In recent years it has become clear that cross-talk between metabolic and immune pathways is central to the regulation of host defence. This interplay appears of particular importance in myeloid cells including dendritic cells and macrophages, but it is unclear how two of their key bacterial sensors NOD2 and TLR2 influence metabolism. Here, we define how NOD2/TLR2 signal in myeloid cells to drive optimal mitochondrial functioning required for bacterial destruction. We uncover a new role for Ataxin-3, a deubiquitinase required for non-selective autophagy, in this pathway. We provide a non-biased analysis of Ataxin-3 targets generating evidence for a role in deubiquitination of metabolic mediators during myeloid cell differentiation that will provide an important basis for further study.

## Introduction

Pattern recognition receptors (PRRs) recognize foreign antigen to direct innate and adaptive immune responses against invading pathogens (1). Polymorphisms in the PRR nucleotide-binding oligomerization domain-containing protein 2 (NOD2) represent the strongest genetic risk factor for the inflammatory bowel disease Crohn’s (CD), and thus this bacterial sensor is the focus of particular research interest (2–4). NOD2 recognizes muramyl dipeptide (MDP), the largest fraction of peptidoglycan, that is present in the cell walls of all bacteria (5). Subsequent activation of NF-κB and MAPK pathways via interaction with receptor-interacting protein kinase 2 (RIPK2) results in an array of immune responses such as production and regulation of pro-inflammatory cytokines (6), and modulation of T-cell function (7–9). NOD2 also directs autophagy, which is important both for bacterial clearance and MHC class II antigen presentation (10). Importantly, NOD2 signaling is intimately linked with that of toll like receptor TLR2, with both responding to ligands derived from the same bacterial component, peptidoglycan. Although the precise mechanisms of cross-regulation are not well understood, both NOD2 and TLR2 activate separate upstream signaling cascades to recruit the same NF-κB and MAPK pathways, and are typically thought to act in a synergistic fashion (11). CD patients harboring NOD2 polymorphisms display loss-of-function for induction of NOD2 and NOD2/TLR2 effector signaling factors (12, 13). In contrast, gain-of-function mutations of NOD2 have been associated with other inflammatory disease such as Blau syndrome and early-onset-sarcoidosis (EOS).

In recent years it has become clear that cross-talk between metabolic and immune pathways is central to the regulation of host defence (14). Immune cells undergo significant metabolic reprogramming during the immune response, both as a result of changes in the metabolic microenvironment induced by inflammation, and in response to immune triggering. This interplay appears of particular importance to dendritic cells and macrophages and controls core processes including differentiation (15). However, while the importance of PRR activation in directing metabolic pathways that impact on immune effector function is now well established, how NOD2 and TLR2 influence myeloid metabolism is unclear. Here, following a phosphoproteomic screen of NOD2 and TLR signaling we identify a deubiquitinase essential for metabolic reprogramming and innate effector function in myeloid cells.

## Results

### NOD2 and TLR2 stimulation leads to ataxin-3 phosphorylation mediated by RIPK2 and TBK1

We identified ataxin-3 as one of the most differentially phosphorylated proteins on NOD2/TLR2 stimulation through a quantitative phosphoproteomic screen in monocyte derived dendritic cells (moDCs) from healthy human donors (Supplementary table 1). Ataxin-3 is a deubiquitinase (DUB) (16) that is required for non-selective autophagy and that is linked to neurodegenerative disease (17–19).

We first validated this result by immunoblotting phospho-enriched samples for ataxin-3 (Figure 1A). Both NOD2 and TLR2 stimulation alone led to ataxin-3 phosphorylation; this effect was enhanced on dual stimulation of NOD2 and TLR2. It was also observed to a lesser extent following stimulation of TLR4, TLR7 and TLR8 (Figure 1A). NOD2 mediated phosphorylation of Ataxin-3 was examined in greater detail. A time course experiment demonstrated that ataxin-3 phosphorylation was maximal 30 minutes after MDP stimulation (Figure 1B). While NOD2 is the only known receptor for MDP, the absolute requirement for NOD2 in the MDP stimulated phosphorylation of ataxin-3 was investigated. We downregulated expression of NOD2 in THP-1 cells using short hairpin RNAs (shRNA) targeting *NOD2* (Figure 1C). Reduction of ataxin-3 phosphorylation on MDP exposure was observed in NOD2 knockdown cells (Figure 1D). Next, given the central importance of RIPK2 in NOD2 signaling (20, 21), the requirement of RIPK2 for phosphorylation of ataxin-3 by NOD2 was investigated. NOD2-RIPK2 inflammatory signalling can be potently and selectively inhibited by the clinically relevant kinase inhibitor Ponatinib, that functions by blocking RIPK2 autophosphorylation and ubiquitination (22). moDCs were treated with Ponatinib prior to stimulation with MDP or PAM_3_CSK_4_ or both, with phosphorylation of p38 used as a positive control for the inhibitor. As expected, inhibition of RIPK2 blocked NOD2 induced phosphorylation of p38, but had no effect on induction by TLR2, which signals to p38 via a MyD88 pathway which is independent of RIPK2 (23). Inhibition of RIPK2 led to complete inhibition of NOD2 induced phosphorylation of ataxin-3, and significant abrogation of the synergistic NOD2/TLR2 signal in both cell types (Figure 1E). Recent evidence suggests that tank binding kinase 1 (TBK1) may represent a novel but important kinase in the NOD2/RIPK2 signalling cascade (24, 25) and MDP stimulation of the NOD2 receptor has been shown to induce TBK1 phosphorylation at S172 (24). Consequently, the requirement for TBK1 in NOD2/RIPK2 dependent phosphorylation of ataxin-3 was examined. We downregulated expression of TBK1 in THP-1 cells using short hairpin RNAs (shRNA) targeting NOD2 (Figure 1F). Reduction of ataxin-3 phosphorylation on MDP exposure was observed in TBK1 knockdown cells (Figure 1G). The possibility that TBK1 might directly phosphorylate ataxin-3, as has been described for a number of other proteins including optineurin (26) and p62 (27), was explored using an *in vitro* kinase assay (Figure 1H). The expected autophosphorylation of TBK1 was demonstrated by a marginally higher molecular weight of the TBK1 band in samples containing both TBK1 and ATP. Importantly, a significant proportion of the ataxin-3 band was noted at a higher molecular weight in samples containing ataxin-3, TBK1 and ATP, consistent with ataxin-3 phosphorylation (Figure 1H). Notably, no change in migration of the ataxin-3 band was seen in samples containing ataxin-3 and TBK1 but not ATP, confirming the ATP dependency of this shift, consistent with phosphorylation. Finally, the phosphorylation site of ataxin-3 was sought, using liquid chromatography mass spectrometry analysis of endogenous ataxin-3 immunoprecipitated from THP-1 cells. A significant shift in mass/charge ratio, consistent with phosphorylation, was detected at a single peptide in the MDP/PAM_3_CSK_4_ stimulated sample only, corresponding to phosphorylation at serine 265 (Figure 1I). This residue has been described as a phosphorylation site in 12 separate large scale mass spectrometry (MS) screens of human primary cells and cell lines (28), and is highly conserved in placental bearing mammals (29), but there is no existing knowledge of its functional relevance. It is located in close proximity to the second ubiquitin interacting motif (UIM), suggesting that phosphorylation could affect specificity of DUB target, as has been described for neighbouring serine residues 256/260/261 (30) (Figure 1J).

**Figure 1.**
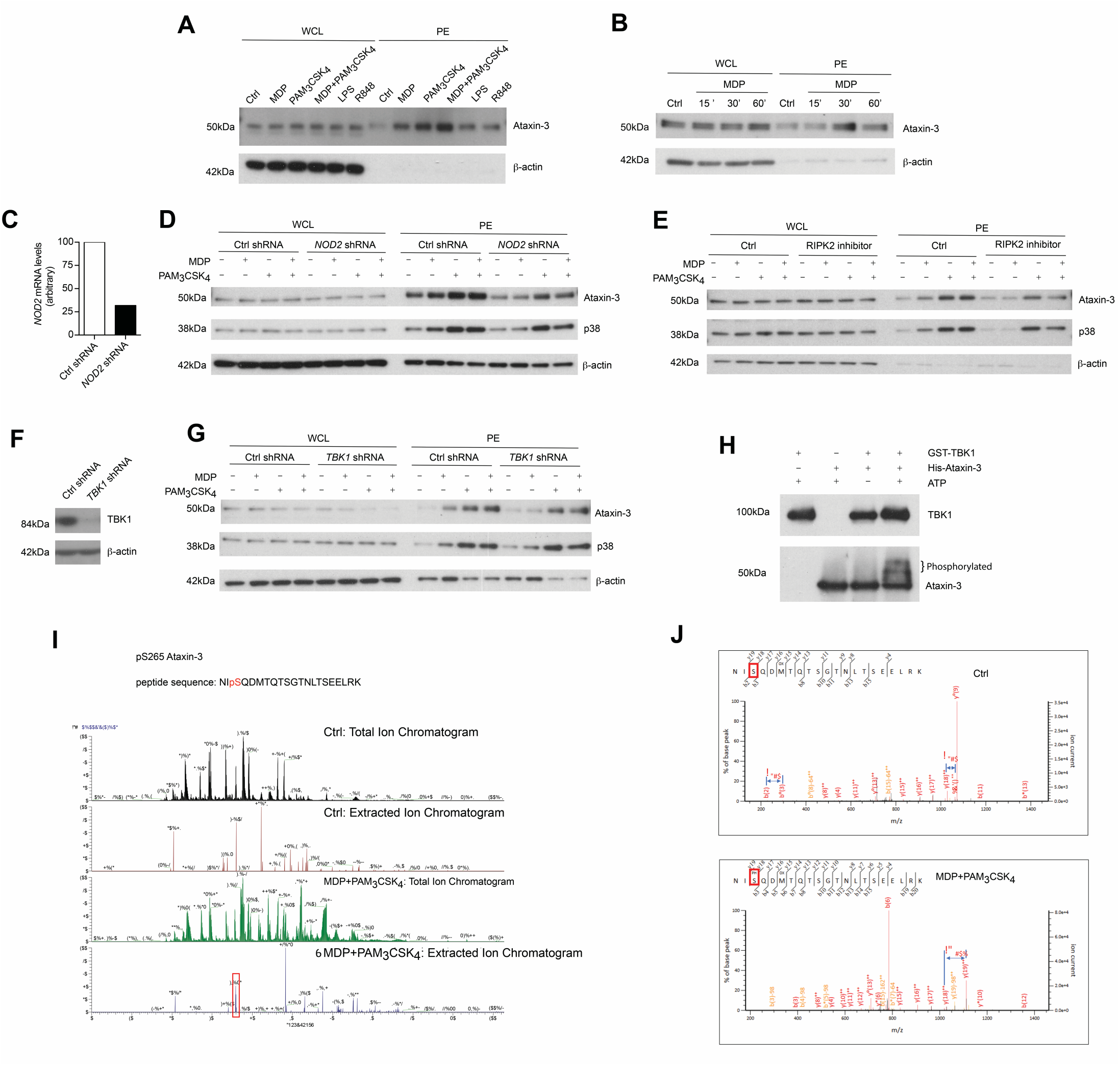
NOD2/TLR2-mediated phosphorylation of ataxin-3. Immunoblot using antibodies against ataxin-3 and β–actin of whole cell and phosphoprotein enriched lysates from moDCs either **(A)** unstimulated or stimulated with the NOD2 ligand MDP 10µg/ml, or the TLR2 ligand PAM_3_CSK_4_ 1µg/ml, or both, or the TLR4 ligand LPS 100ng/ml or the TLR7/8 ligand R848 (Resiquimod) 1µg/ml for 30 minutes or **(B)** unstimulated or stimulated with the NOD2 ligand MDP for 15, 30 or 60 minutes. (**C**) THP-1 cells were transduced with control or *NOD2*-targeting lentiviral shRNAs and analysed for *NOD2* expression by qPCR analysis. **(D)** Immunoblot using antibodies against ataxin-3, p38 and β–actin of whole cell and phosphoprotein enriched lysates from THP-1 cells expressing control or NOD2 shRNA and either unstimulated or stimulated with the NOD2 ligand MDP or the TLR ligand PAM_3_CSK_4_, or both, for 60 minutes. **(E)** Immunoblot using antibodies against ataxin-3, p38 and β–actin of whole cell and phosphoprotein enriched lysates from THP1 cells which were pre-treated with the RIPK2 inhibitor Ponatinib 50nM for 60 minutes and then left unstimulated or stimulated with the NOD2 ligand MDP or the TLR2 ligand PAM_3_CSK_4_ or both. (**F**) THP-1 cells were transduced with control or *TBK1*-targeting lentiviral shRNAs and immunoblotted to detect TBK1 expression **(G)** Immunoblot using antibodies against ataxin-3, p38 and b-actin of whole cell and phosphoprotein enriched lysates from THP-1 cells expressing control or TBK1 shRNA left unstimulated or stimulated with the NOD2 ligand MDP or the TLR2 ligand PAM_3_CSK_4_ or both for 60 minutes. **(H)** Immunoblot using antibodies against TBK1 and ataxin-3 following an *in vitro* kinase assay of GST-TBK1 protein or His-ataxin-3 protein with ATP, or both GST-TBK1 and His-ataxin-3 with or without ATP which were incubated for 60 minutes at 30° C. **(I)** Identification of the phosphorylated serine residue (S265) with a characteristic increase in mass/charge ratio in the stimulated sample. **(J)** A 3-dimensional computer model of the structure of ataxin-3, with the S265 phosphorylation site indicated. All immunoblots are representative of at least two independent experiments.

Taken together, this data shows that activation of NOD2/TLR2 signaling pathway induces phosphorylation of the DUB ataxin-3. TBK1 is required for the direct phosphorylation of ataxin-3 at serine 265 following NOD2/TLR2 activation.

### Ataxin-3 localizes with the mitochondrial cristae protein MIC60 and regulates the expression of the oxphos machinery components

To identify novel interacting partners of ataxin-3 in innate immune cells, endogenous ataxin-3 was immunoprecipitated in moDCs from healthy human donors and subjected to mass spectrometry analysis. One of the most abundant proteins identified in the pull down was the mitochondrial cristae protein MIC60 (Figure 2 A, B). The association between ataxin-3 and MIC60 was validated through immunoblot of immunoprecipitated ataxin-3 (Figure 2C). There appeared to be no change in abundance of MIC60 when moDCs were stimulated with MDP + PAM_3_CSK_4_ prior to immunoprecipitation of ataxin-3, suggesting that NOD2/TLR2 stimulation does not affect binding. To further confirm the association, MIC60 was pulled down, and an immunoblot for ataxin-3 performed (Figure 2D). Here, ataxin-3 appeared at two molecular weights, suggesting that either two separate isoforms bind to MIC60, or it is post-translationally modified. Finally, super-resolution assessment using stimulated emission depletion (STED) microscopy confirmed that ataxin-3 was situated in close proximity to MIC60 (Figure 2E).

**Figure 2.**
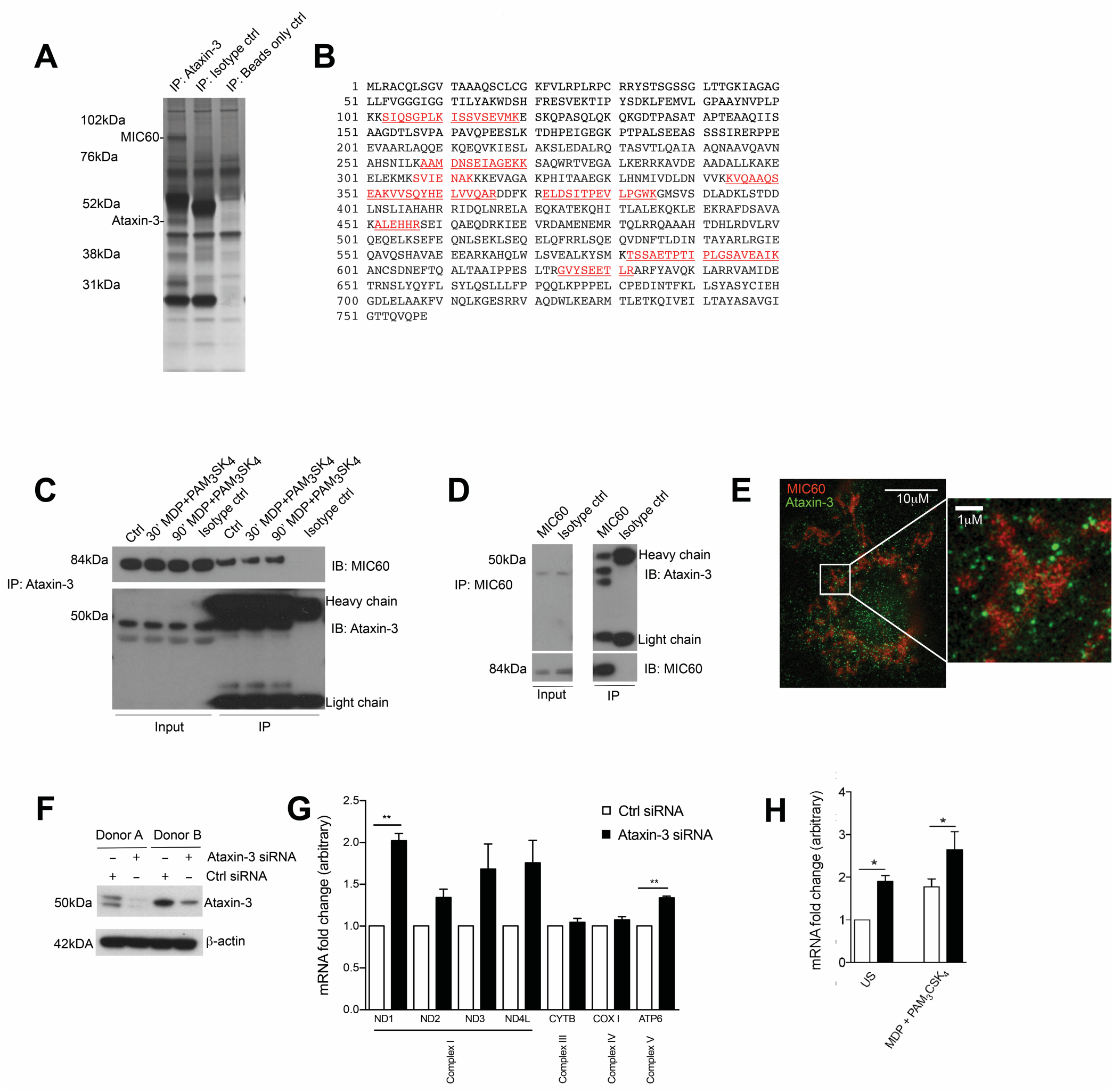
Ataxin-3 associates with the mithocondrial MIC60. **(A)** Silver stain of immunoprecipitation of ataxin-3 in moDCs, using ataxin-3 antibody, isotype control antibody or immunoprecipitation beads only. **(B)** The sequence coverage of MIC60 identified following mass spectrometry analysis of the immunoprecipitation of ataxin-3 in moDCs, with the peptides in red indicating the identified peptides. **(C)** Immunoblot using antibodies against MIC60 and ataxin-3 of lysates where ataxin-3 has been immunoprecipitated from moDCs either left unstimulated or stimulated for 30 or 60 minutes with MDP+PAM3CSK4. **(D)** Immunoblot using antibodies against MIC60 and ataxin-3 of lysates where MIC60 has been immunoprecipitated from moDCs. **(E)** STED microscopy using antibodies against ataxin-3 and MIC60 in moDCs. **(F)** Immunoblot using antibodies against ataxin-3 and β–actin in moDCs following transfection with control or ataxin-3 siRNA. **(G)** RT-qPCR analysis of selected mitochondrial genome encoded transcripts following ataxin-3 depletion by siRNA in moDCs; n=3, one way ANOVA **p<0.01 **(H)** RT-qPCR analysis of MT-ND1 mRNA expression following ataxin-3 depletion by siRNA in moDCs subsequently left unstimulated or stimulated for 6 hours with MDP+PAM_3_CSK_4_; n=3, paired t-test *p<0.05,

MIC60 is the largest protein in the mitochondrial contact site (MICOS) complex, which is embedded in the mitochondrial inner membrane and acts as a key regulator of cristae junction formation and assembly of respiratory chain complexes which are required for oxidative phosphorylation (31). Additional specific roles for MIC60 include the import of proteins (32, 33) and regulation of mitochondrial DNA (mtDNA) transcription (34, 35). Consequently, we examined the effect of ataxin-3 knockdown in moDCS on expression of mtDNA genes that encode components of the oxphos machinery. moDCS from healthy human donors were found to express either a single or double isoform of ataxin-3 with approximately equal frequency, but all detectable isoforms could be efficiently knocked down (Figure 2F). Ataxin-3 depletion led to a two-fold upregulation of NADH-ubiquinone oxidoreductase 1 (MT-ND1), with a statistically significant upregulation observed for mRNA expression of two other genes from Complex I, MT-ND3 and MT-ND4L (Figure 2G). In comparison, genes encoding components of Complex II, III and IV were broadly unaffected. It is noteworthy that mtDNA transcription is tightly regulated due to its close links to oxphos, and thus 1.5 to 2 fold changes in mRNA expression level represent a potentially functionally significant alteration (36). The effect of NOD2/TLR2 stimulation was examined on MT-ND1, the most significantly affected gene. Stimulation led to a significant upregulation in mRNA expression in both the control and ataxin-3 depleted cells, but there was significantly greater ND1 in the ataxin-3 depleted cells following stimulation (Figure 2H).

These results indicate a novel association between ataxin-3 and MIC60, a component of the MICOS complex involved in the regulation of mtDNA. We demonstrated that ataxin-3 regulates mtDNA by downregulating the expression of Complex I genes, an effect increased on NOD2/TLR2 sensing.

### Ataxin-3 is important for optimal mitochondrial respiration following NOD2 and TLR2 stimulation

We next investigated the functional relevance of the observed changes in the expression of Complex I genes. Using short hairpin RNAs (shRNA) targeting *ataxin-3*, we downregulated ataxin-3 expression in THP-1 cells (Figure 3a). We next performed a real time analysis of oxidative phosphorylation to address the function of Ataxin-3. We found that Ataxin-3 depletion led to a significant reduction in all the key parameters of mitochondrial respiration assessed (Figure 3B-F).

**Figure 3.**
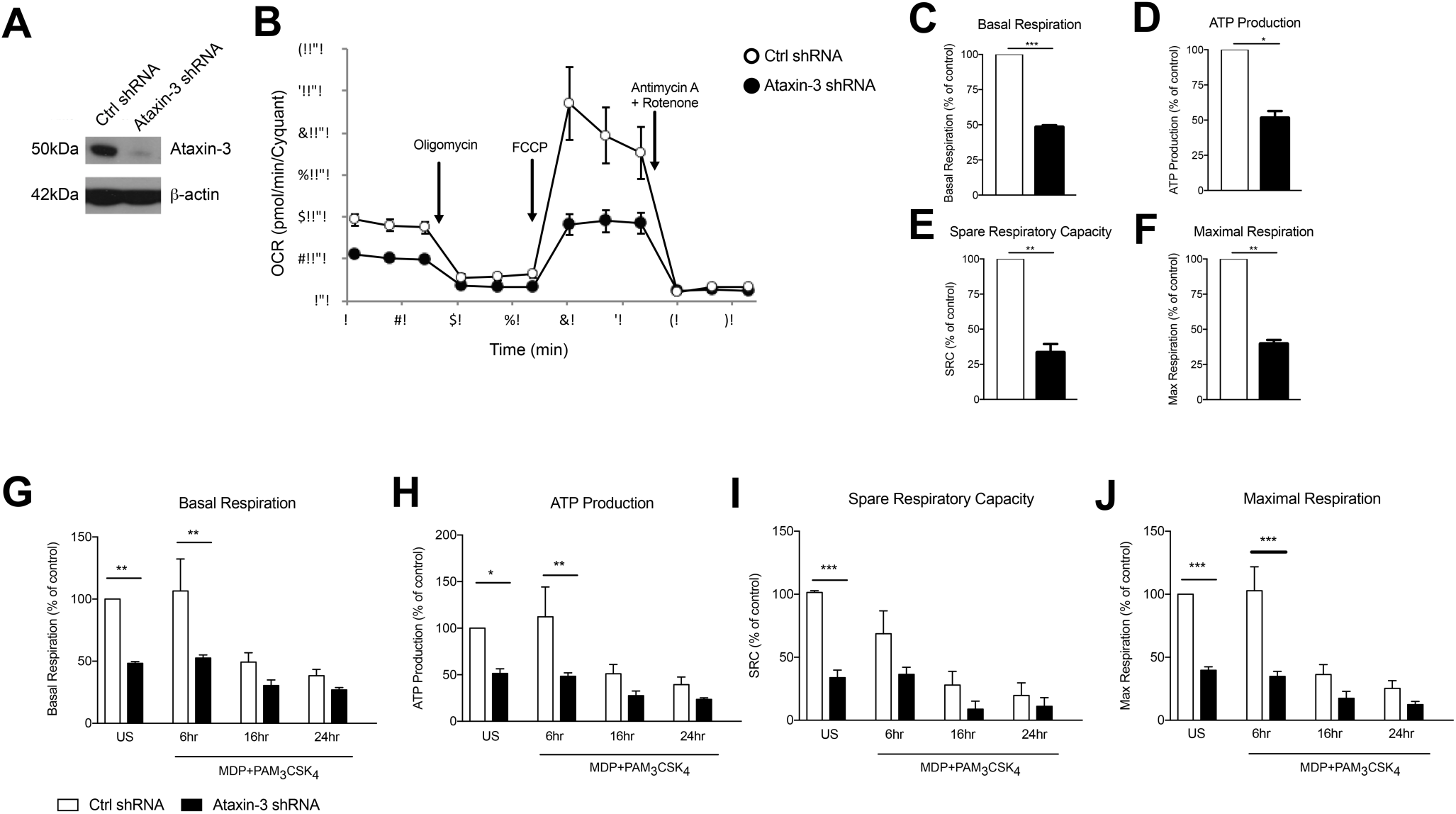
Ataxin-3 is required for mitochondrial respiration in immune cells. **(A)** Immunoblot using antibodies against ataxin-3 and β–actin in THP1 cells following transfection with control or ataxin-3 shRNA. **(B)** Representative Seahorse sequential OCR measurements by Seahorse of control and ataxin-3 shRNA THP1 cells. **(C-F)** Seahorse analysis of key mitochondrial respiration parameters in control shRNA cells and ataxin-3 shRNA THP1 cells. **(G-J)** Seahorse analysis of key mitochondrial respiration parameters in control shRNA cells and ataxin-3 shRNA THP1 cells either left unstimulated or stimulated for 6 hours, 16 hours or 24 hours with MDP+PAM3CSK4. (C-J) n=3, paired t-test *p<0.05, **p<0.01, ***p<0.001. All OCR measurements were normalised to cell number. (O-R) n=3, one way ANOVA *p<0.05, **p<0.01, ***p<0.001

We next determined whether NOD2/TLR2-mediated ataxin-3 phosphorylation stimulation affects mitochondrial respiration. Prolonged triggering of these PRRs led to an expected metabolic shift (37) with downregulation of oxidative phosphorylation. In the ataxin-3 depleted cells the level of oxphos remained significantly lower than in the control cells following stimulation (Figure 3G-J). Importantly, no significant difference in mtDNA copy number was found between the control and ataxin-3 depleted THP-1 cell line (Supplementary figure 1), suggesting that there are no differences in mitochondrial mass or turnover through mitophagy to explain the observed changes.

Taken together, this data shows that in innate immune cells, ataxin-3 is required for optimal mitochondrial respiration and this effect is enhanced following its phosphorylation on NOD2/TLR2 stimulation.

### Ataxin-3 is required for mitochondrial ROS production, and is necessary for optimal bacterial killing

A key function of mitochondrial respiration in immune cells is the generation of mROS. This results from leakage of electrons, predominantly from Complex I and to a lesser extent from Complex III, which partially reduce oxygen to form superoxide (38). The effect of ataxin-3 depletion on mROS and total cellular ROS was therefore examined. Ataxin-3 depletion led to a significant reduction in mROS (Figure 4A). As expected, there was a corresponding decrease in total cellular ROS, to which mROS makes a significant contribution (Figure 4B). The ability of immune cells to upregulate mROS production on pathogen challenge is crucial to the immune response (39); importantly, ataxin-3 depleted cells also produced less mROS on NOD2/TLR2 stimulation (Figure 4C). mROS forms an important component of antibacterial responses, and is important for bacterial killing (39). To test the functional significance of the observed impairment of mitochondrial oxphos and mROS generation on NOD2/TLR2 triggering, we assessed the response of ataxin-3 depleted macrophages to *Salmonella typhimurium*, a Gram-negative intracellular bacterium that is sensitive to ROS-dependent killing (39, 40). A gentamicin survival assay was undertaken. While bacterial invasion was unchanged, as evidenced by similar bacterial counts at one hour post infection, there was subsequently significantly greater bacterial survival in the ataxin-3 depleted cells that was maximal at six hours (Figure 4D). There were no significant differences in cell viability between the two groups at any of the infection time points (Supplementary figure 2), excluding differences in cell survival as a contributing factor in the bacterial killing deficit.

**Figure 4.**
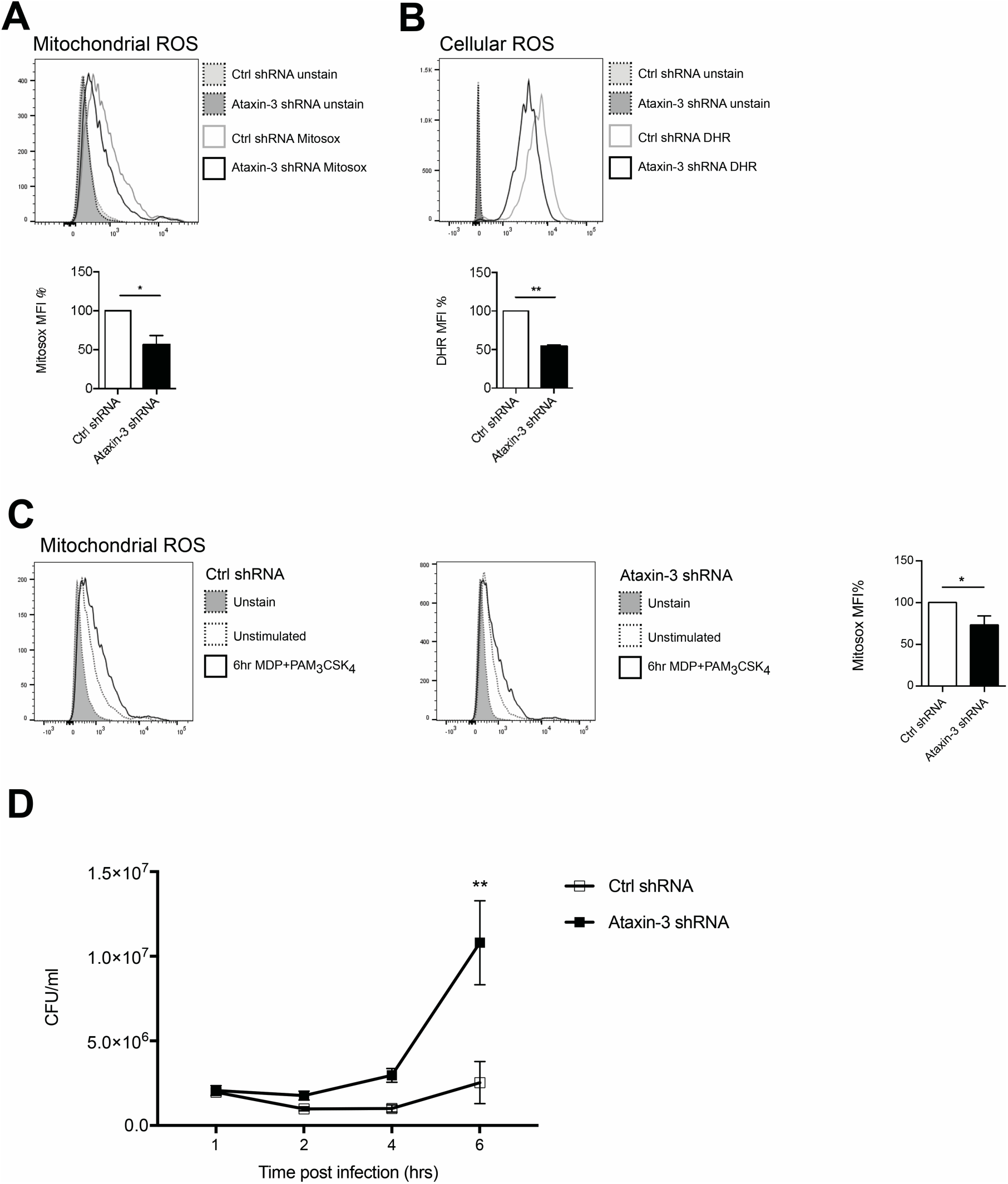
Ataxin-3 is required for ROS production and bacterial killing. **(A)** FACS analysis of Mitosox to quantify mROS; n=3 Student’s paired t test, * p<0.05. **(B)** FACS analysis of Mitosox of DHR to quantify total cellular ROS in control and ataxin-3 shRNA THP-1 cells. n=7 Student’s paired t test, **p<0.01. **(C)** Mitosox to quantify mROS in control and ataxin-3 shRNA THP-1 cells either left unstimulated or following stimulation for 6 hours with MDP+PAM_3_CSK_4_; n=4, Student’s paired t test, * p<0.05. (**D**) Gentamicin survival assay of *Salmonella typhimurium* infected ataxin-3 shRNA THP-1 cells compared to control. Student’s paired t test, **p<0.01.

Taken together, this data shows that in macrophages ataxin-3 is required for mROS generation and this contributes to effective intracellular bacterial killing.

### An unbiased ubiquitome screen reveals novel DUB targets of ataxin-3 following NOD2 and TLR2 activation

To further define the role of ataxin-3 in NOD2/TLR2 signaling, we sought to define downstream DUB targets. Classically the ubiquitinated proteome, also termed the ubiquitome, was first enriched using His_6_-tagged ubiquitinated conjugates under denaturing conditions. However, concerns exist over the impact of overexpressed modified His_6,_ which competes with endogenous ubiquitin, on the ubiquitome (41). An alternative method of enrichment, using high affinity ubiquitin traps has therefore been developed. These tandem ubiquitin binding entities (TUBEs) specifically recognise, bind to, and stabilise polyubiquitinated proteins, protecting them from degradation by DUBs or the proteasome. The enriched ubiquitinated proteins can then be analysed by MS (41–43). The ability of TUBEs to preferentially recognise either K48 or K63 linked polyubiquitin provides a further advantage. Consequently, the use of a K63-TUBEs1 system which shows a 10 fold higher affinity for K63 linked chains, allows the selective enrichment of the K63 linked ubiquitome. This provides a specific means of purifying the K63 chains favoured by ataxin-3 (44), and thus was employed as a strategy for defining novel DUB targets of ataxin-3 in immune cells The TUBEs2 system was used to enrich ubiquitinated proteins in ataxin-3 depleted and control THP-1 cells, either left unstimulated or stimulated for one hour with MDP + PAM_3_CSK_4_. Samples from 3 biological replicates were then subjected to mass spectrometry analysis (Figure 5A). As a quality control prior to MS analysis, immunoblotting with an antibody against K63-linkage specific polyubiquitin demonstrated a marked increase in the levels of K63 ubiquitinated proteins in the ataxin-3 depleted cells, most notably at higher molecular weights, with NOD2/TLR2 stimulation leading to a separate shift in staining pattern (Figure 5B).

**Figure 5.**
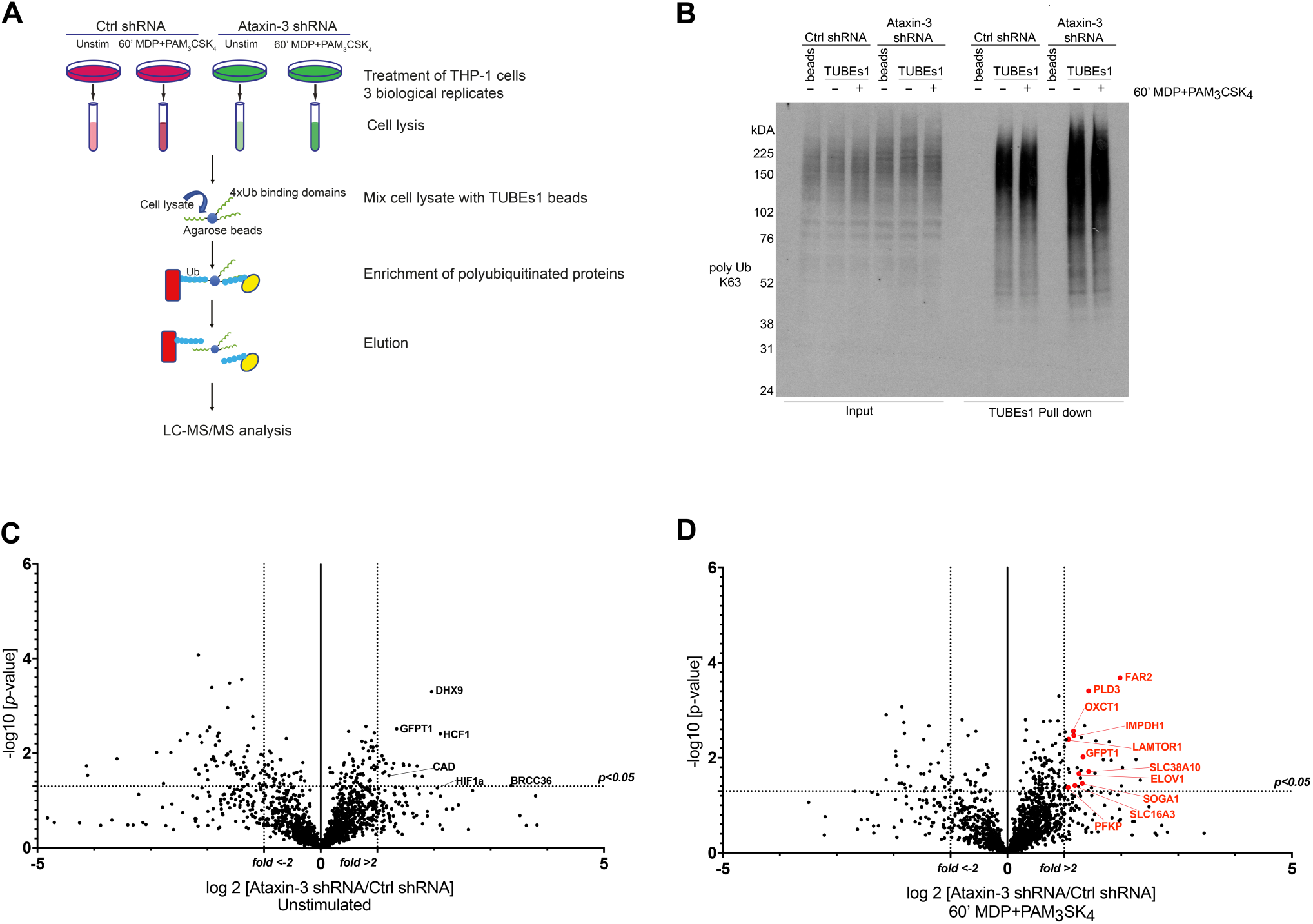
Ubiquitome screen reveals novel DUB targets of ataxin-3. **(A)** Flow chart showing experimental protocol. Control and ataxin-3 shRNA THP-1 cells were left unstimulated or stimulated for 60 minutes with MDP+PAM3CSK4 prior to ubiquitin enrichment using TUBEs1 beads, with the lysates subsequently subjected to LC-MS/MS analysis. **(B)** Immunoblot using antibody against K63 linked polyubiquitinated proteins of whole cell lysates (input) and TUBEs1 enriched fractions or beads only control samples. Volcano plots comparing ubiquitinated protein expression in **(C)** unstimulated control and ataxin-3 shRNA THP-1 cells and **(D)** control and ataxin-3 shRNA THP-1 cells following MDP+PAM3CSK4 stimulation. The dotted lines on the y-axes represents a p value of 0.05 (t-test), and on the x-axes represents fold change >2 or <-2.

Mass spectrometry analysis identified 291 proteins as changing significantly between any condition when averaged across the three biological replicates. As ataxin-3 acts as a DUB, ataxin-3 depletion would classically lead to an accumulation of ubiquitinated targets and thus particular attention was paid to those proteins that showed an increase in abundance in the ataxin-3 depleted samples (table 2). However, as deubiquitination can also regulate protein stability, it is likely that a number of the proteins found to decrease in abundance on ataxin-3 depletion are also direct DUB targets of ataxin-3. Most notably, the immunometabolic regulator HIF1α (45) was found to be more abundant in the ataxin-3 depleted cells, suggesting that ataxin-3 may deubiquitinate HIF1α (Figure 5C). To specifically interrogate the importance of NOD2/TLR2 phosphorylation of ataxin-3 on DUB activity, the abundance of proteins in the ataxin-3 depleted cells was compared to the control cells following stimulation with MDP+PAM_3_CSK_4_ (Supplementary table 3). Strikingly, a cluster of proteins related to metabolism were noted in the ataxin-3 depleted cells (marked in red on Figure 5D).

### Ataxin-3 deubiquitinates HIF, PLD3 and LAMTOR1 upon NOD2 and TLR2 activation

A number of the proteins from the mass spectrometry analysis were selected for further validation. HIF1α was of particular interest given its central role in immunometabolism. Although classically described as part of the family of hypoxia-inducible factor regulators, mediating the cellular response to hypoxia (46), more recent work has demonstrated a broader role in regulation of the immune system. It is important for both the survival and function of cells of the innate immune system, through regulation of metabolic activation (47–49). HIF1αwas validated as a DUB target of ataxin-3 through immunoblot, with a significant increase in ubiquitinated HIF1α in the ataxin-3 depleted cells (Figure 6A). HIF1α was detected in the whole cell lysates of both control and ataxin-3 depleted cells in normoxia, as the proteasome inhibitor MG132 which reduces the degradation of all ubiquitinated proteins was added to the cell suspension 30 minutes before the end of all TUBEs experiments.

**Figure 6.**
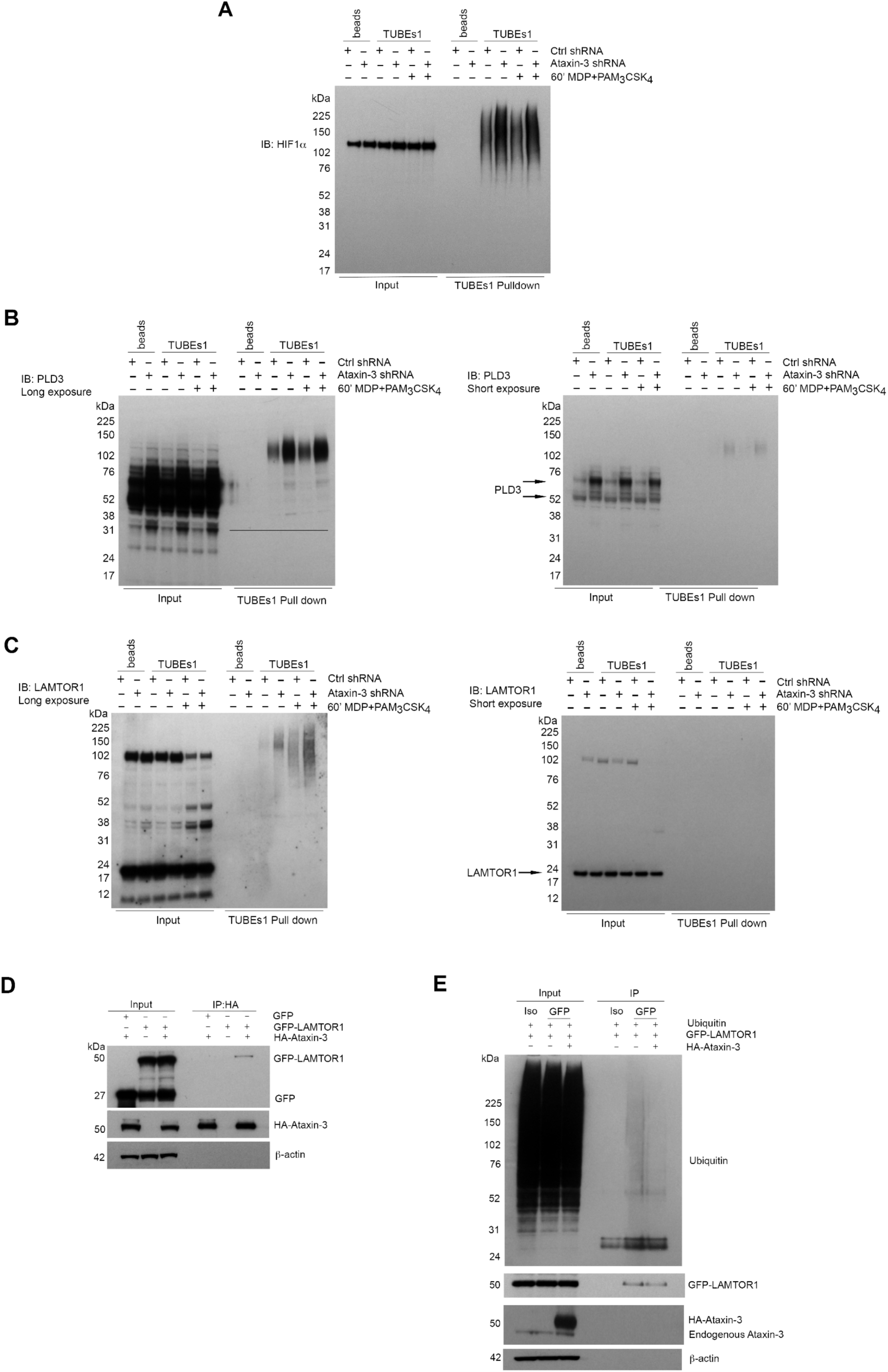
Validation of novel DUB targets of ataxin-3. **(A)** Immunoblot using antibody against HIF1a in whole cell lysate (input) and TUBEs1 enriched fractions from control and ataxin-3 shRNA THP-1 cells left unstimulated or stimulated with 60’ MDP+PAM3CSK4. Immunoblot using antibodies against **(B)** PLD3 and **(C)** LAMTOR1 in whole cell lysates (input) and TUBEs1 enriched fractions from control shRNA and ataxin-3 shRNA THP-1 cells left unstimulated or stimulated for 60 minutes with MDP+PAM3CSK4. **(D)** Immunoblot using antibodies against GFP, HA and β–actin in input and HA-immunoprecipitated lysates, in HEK293 cells where GFP and/or GFP-LAMTOR1 and/or HA-ataxin-3 were overexpressed. **(E)** Immunoblot using antibodies against ubiquitin, GFP, ataxin-3 and β–actin in input and isotype control (iso) or GFP immunoprecipitated lysates where ubiquitin and GFP-LAMTOR1 were overexpressed with or without HA-ataxin-3.

Phospholipase D3 (PLD3) was also intriguing as it has recently been shown to regulate inflammatory cytokine responses in response to TLR9 signaling by acting as an endonuclease (50). In humans, mutations confer an increased risk for the neurodegenerative diseases Alzheimer’s (51) and spinocerebellar ataxia(52), with increasing evidence of the role of innate immune dysfunction in neurodegeneration. (53). Importantly, polyglutamine repeat mutations in ataxin-3 are themselves associated with the spinocerebellar ataxia Machado-Joseph disease (18). Ragulator complex protein LAMTOR1 forms part of the Ragulator complex essential for amino acid sensing and activation of mTORC1 (54), but has also been linked to independent roles in lysosomal maturation (55) and M2 macrophage differentiation (56). PLD3 and LAMTOR1 were both validated by immunoblot (Figure 6B and C). Ataxin-3 depletion led to accumulation of ubiquitinated forms of both PLD3 and LAMTOR1. Notably, NOD2/TLR2 stimulation increased LAMTOR1 ubiquitination in the control cells, but there was markedly more LAMTOR1 ubiquitination in the stimulated ataxin-3 depleted cells (Figure 6C). This suggests that ataxin-3 modulates the ubiquitination of LAMTOR1 induced by NOD2/TLR2 stimulation.

To further study the interaction between LAMTOR1 and ataxin-3, an overexpression system was employed in the human cell line HEK293. HA-ataxin-3 and GFP-LAMTOR1 was overexpressed, together with a GFP-control and then IP of HA-ataxin-3 performed using an antibody against the HA tag. Ataxin-3 was found to bind directly to LAMTOR1 (Figure 6D). Next, the finding that LAMTOR1 is a direct DUB target of ataxin-3 was confirmed. Ubiquitin, GFP-LAMTOR1 and HA-ataxin-3 were overexpressed in HEK293 cells and IP of GFP-LAMTOR1 performed using an antibody against the GFP tag. The level of ubiquitinated GFP-LAMTOR1 was found to be significantly less in cells where HA-ataxin-3 was co-expressed (Figure 5E).

Taken together, these results show that ataxin-3 modulates the ubiquitination of previously unidentified targets related to metabolism. This effect is enhanced on following phosphorylation of ataxin-3 on NOD2/TLR2 triggering.

## Discussion

In this study we demonstrate that NOD2 and TLR2 phosphorylates the deubiquitinase ataxin-3 at serine 265 through a signaling cascade involving RIPK2 and TBK1. Immunoprecipitation and MS analysis of interacting partners established an association with a core component of the mitochondrial MICOS complex MIC60. Ataxin-3 was subsequently shown to be necessary for optimal mitochondrial respiration and mitochondrial ROS generation in macrophages, an effect enhanced by its phosphorylation on NOD2/TLR2 triggering. In line with this, we found that ataxin-3 is required for optimal intracellular bacterial killing of *Salmonella typhimurium*. Finally, we dissected the specific DUB role of ataxin-3 in an immune context through an unbiased MS screen of the ubiquitome. A preponderance of metabolism related proteins were discovered including HIF1α, phospholipase D3 and LAMTOR1, underlining a central role of ataxin-3 in immunometabolism.

Deubiquitinating enzymes (DUBs) represent specialised proteases which modify ubiquitin chains by cleaving the isopeptide bonds linking the ubiquitin C-terminus to a lysine side chain on the target protein. Modification of ubiquitination may alter cellular responses through regulation of target protein stability, or mediate signal transduction through non-degradative pathways including mediation of protein-protein interactions (57). Ataxin-3 is a small protein, consisting of 364 amino acids, that is ubiquitously expressed (58). At the N-terminus there is a catalytic Josephin domain, which acts as a protease that hydrolyses ubiquitin linkages and allows ataxin-3 to function as a DUB (59). A flexible C-terminal tail contains either two or three ubiquitin-interacting motifs (UIMs), according to the isoform (60). The UIMs mediate selective binding to ubiquitin chains, determining the type of chain that can be cleaved by the Josephin domain. Ataxin-3 shows a preference for cleavage of K63-Ub chains, although it is able to bind both K63 and K48 chains (44).

Ataxin-3 has been linked to neurodegenerative disease after unstable CAG repeat expansions in the ATXN3 gene were identified as the cause of spinocerebellar ataxia Type 3 (SCA3), also known as Machado-Joseph Disease, the most common autosomal dominant ataxia (61). Importantly, the expanded polyglutamine stretch results in more complex sequelae than a simple loss of protein function, and likely leads to a toxic gain of function through altered binding properties, aggregation and subcellular localisation (62). Accumulating evidence suggests that ataxin-3 performs diverse cellular roles, including DNA repair (63, 64) and transcriptional regulation (65), regulation of protein quality through endoplasmic reticulum (ER) associated degradation (66) and aggresome formation (67), and beclin-1 dependent autophagy (17). A recent study provided a first link to immune regulation, demonstrating that ataxin-3 regulates Type 1 interferon antiviral responses through interaction with histone deacetylase 3 (HDAC 3) (68). The present study is the first to link ataxin-3 to PRR signaling and demonstrate its importance in mitochondrial respiration in macrophages, following the discovery that ataxin-3 associates with the mitochondrial protein MIC60.

MIC60 forms a core part of the MICOS complex, which is embedded in the mitochondrial inner membrane. It acts as a key regulator of mitochondrial inner membrane shape and organisation. This is essential for cristae junction formation and assembly of respiratory chain complexes which are required for oxidative phosphorylation (31). MIC60 also appears to act independently of the MICOS complex, and has recently been implicated in regulation of mtDNA transcription (35). The mitochondrial genome encodes just 13 proteins, all essential components of oxphos complexes I, III and IV. In keeping with this known function of MIC60, we found that depletion of ataxin-3 led to specific upregulation of mtDNA transcripts encoding proteins required for Complex I, and this was further upregulated by NOD2/TLR2 stimulation. In addition, specific interrogation of mitochondrial oxidative phosphorylation through use of the Seahorse platform demonstrated that ataxin-3 depletion led to a particularly striking impairment in maximal respiration and spare respiratory capacity (SRC). SRC represents the extra mitochondrial capacity available within a cell to produce energy under conditions of increased work or stress and is important for cellular function and survival (69–71). Macrophages increase their SRC in response to bacterial infection to drive anti-microbial responses, and this is coordinated in part by modulation of the ETC Complexes I and II (72). In M2 macrophages, SRC is critical for their activation and prolonged survival, and clearance of the parasitic helminth (73). We found that ataxin-3 depletion impaired mROS production at both baseline and in response to NOD2/TLR2 stimulation, demonstrating the functional relevance of the observed mitochondrial respiratory impairment. Finally, we demonstrated the importance of ataxin-3 in intracellular killing of the pathogen *Salmonella typhimurium*, with PRR mediated mROS generation well established as critical for destruction of this bacterium (39).

Fewer than 100 DUBs are thought to be responsible for regulating the ubiquitination of tens of thousands of proteins in a tightly regulated and sophisticated manner (57, 74). Hence defining the wide-ranging DUB targets of ataxin-3 is essential to decipher its functions. Ubiquitin signalling represents an indispensable mechanism of regulating both the innate and adaptive immune response, and is central to the NOD2 cascade (75). For the first time, this study undertook an unbiased screen of the ubiquitome in ataxin-3 depleted cells. Notably, a significant number of metabolism related proteins were found, further reinforcing the potential importance of ataxin-3 in cellular immunometabolism.

The discovery that HIF1α is a DUB target of ataxin-3 in macrophages is noteworthy, given the emergency of HIF1α as a key immunometabolic regulator. Work in a conditional HIF1 α knockout mouse, targeting the myeloid lineage, demonstrated the critical requirement of HIF1 α for inflammatory responses (76). This correlated with defects in glycolysis and metabolic activation, which is tightly regulated by HIF1 α The endolysosomal protein PLD3 was also validated as a further novel DUB target of ataxin-3. Although PLD3 contains two phosphodiesterase domains, and hence is classed as a member of the phospholipase diesterase (PLD) family which act to hydrolyse phospholipids, the phospholipase activity of PLD3 has not been definitively established. The importance of PLDS in immunity was recently demonstrated, with the discovery that it acts as a single-stranded acid exonuclease that breaks down ligands for the PRR TLR9, hence regulating TLR9 mediated inflammatory responses in collaboration with PLD4 (50).

The discovery in the present study that NOD2/TLR2 triggering leads to deubiquitination of LAMTOR1 by ataxin-3 is of particular interest. LAMTOR1, also known as p18, is a late endosome/lysosome membrane adapter protein that localises to the lipid rafts of these organelles (77, 78). LAMTOR1 plays an essential role in the activation of the mTORC kinase complex in response to amino acid levels (54). Through a mechanism involving the lysosomal v-ATPase in the presence of amino acid sufficiency, LAMTOR1 forms a scaffold at the lysosomal membrane with LAMTOR2,3,4 and 5 (the pentameric Ragulator complex) for the Rag GTPase complex (RagAB/CD) (54, 79, 80). This leads to the recruitment and activation of mTORC1 which inhibits autophagy. The lysosomal v-ATPAse-Ragulator complex also activates another critical metabolic sensor, AMP-activated protein kinase (AMPK), which responds to falling energy levels by driving cellular catabolism programmes and downregulating anabolic pathways(81). Thus this complex is able to respond to both energy/nutrient sufficiency and deficiency. Strikingly, of all cells, macrophages express the highest levels of the five Ragulator components, suggesting their importance in the immune response (82). Indeed, LAMTOR1 was recently found to be essential for the polarisation of M2 macrophages both *in vitro* and *in vivo* in a knockout mouse model, by coupling metabolism to immunity (56).

The physiological importance of NOD2 and TLR2 in both the innate and adaptive immune response is well established. The interplay between NOD2 and TLR2 has been well characterized given the fact that they both respond to adjacent components of PGN found in the bacterial cell walls. The NOD2 signaling pathway amplifies TLR2 activation and both receptors synergize in the induction of cytokine production. *NOD2* variants confer the greatest single genetic risk factor for Crohn’s disease disease (2, 3), yet significant gaps remain in our knowledge of how this receptor exerts its effects (83). Notably, despite the recent explosion of interest in the field of immunometabolism, almost nothing is known about how the synergistic effects of NOD2 and TLR2 signaling might intersect with metabolic pathways to modulate the immune response. Here, we defined the molecular and functional basis by which NOD2/TLR2 sensing links to ataxin-3 and, consequently, other immunometabolic factors. Future studies are required to provide novel prospects for modulating these pathways as new therapeutic strategies for inflammatory disorders.

## Materials and Methods

### Cells

Human monocytes were purified from healthy donor peripheral blood mononuclear cells (PBMCs) by positive immunoselection with anti-CD14-conjugated MACS beads (Miltenyi Biotec). moDCs were obtained by culturing monocytes for 5 days with IL-4 and GM-CSF (Peprotech). Immature moDCs were harvested on day 5 of culture. The human THP-1 cell line was purchased from ATCC. Prior to use, THP-1 were differentiated by treatment with 25ng/ml phorbol 12-myristate 13-acetate (PMA) (Sigma) for 16 hr.

### Reagents and antibodies

The following stains were used: MItosox Red M36008 (Invitrogen) and DHR 123 D23806 (Invitrogen). Antibodies include mouse anti-human ataxin-3 65042 1H9-2 (BioLegend), mouse anti-human MIC60 ab110329 (Abcam), mouse anti-human MIC60 ab137057 (Abcam), rabbit anti-human TBK1 #3504 D1B4 (Cell Signaling), rabbit anti-human RIPK2 #4142 D10B11 (Cell Signaling), rabbit anti-human p38 #9212 (Cell Signaling), rabbit anti-human LAMTOR1 #8975 D11H6 (Cell Signaling), rabbit anti-human PLD3 HPA012800 (Sigma), rabbit anti-human HA #3724 C29F4 (Cell Signaling). The secondary antibodies included: anti-rabbit HRP conjugate #7074 (Cell Signaling), anti-mouse HRP conjugate #7076 (Cell Signaling), goat anti-rabbit Alexa fluor 488 A-11034 (Invitrogen), goat anti-rabbit Alexa fluor 488 A-11029 (Invitrogen), goat anti-mouse Alexa fluor 568 A-11036 (Invitrogen), goat anti-rabbit Alexa fluor 568 A-11004 (Invitrogen). Beta actin HRP conjugate #5125 (Cell Signaling). For qPCR, the following Taqman primers were used (all ThermoFisher): NOD2 (Hs01550753_m1), RPLP0 (Hs99999902_m1), MT-ND1 Hs02596873_s1, MT-ND2, Hs02596874_g1, MT-ND3 Hs02596875_s1, MT-ND4L Hs02596877_g1, MT-CYB Hs02596867_s1, MT-CO1 Hs02596864_g1, ATP6, Hs02596862_g1. For overexpression experiments, GFP-LAMTOR1 and HA-ataxin-3 were obtained from the University of Dundee.

### Cell stimulation

moDCs and THP-1 cells were left unstimulated or stimulated with 10 µg/ml MDP or 1 µg/ml PAM_3_CSK_4_ (Invivogen) or both at the indicated time points. In some experiments, other PRR ligands were used including LPS 100 ng/ml, and R848 1 µg/ml (Invivogen) or cells were treated with the small molecule inhibitor Ponatinib (50nM) for 1 hr.

### Phosphoprotein purification

Cells were harvested on ice and washed once with ice cold modified Hanks Buffered Saline (HBS) (20mM HEPES pH 7.4, 150mM NaCl in ddH_2_0). Cell pellets were lysed in Qiagen “Phosphoprotein lysis buffer” containing 0.25% (v/v) CHAPS with 1% (v/v) phosphatase inhibitor cocktail 3 (Sigma), protease inhibitor tablet (Qiagen) and the nuclease 0.0002% (v/v) Benzonase (Qiagen) at 4°C for 40 minutes, with vortexing every 10 minutes. Cell debris was removed by centrifugation of the lysate at 13300rpm for 30 minutes at 4°C. The clarified supernatant was then transferred to fresh pre-cooled tubes and protein concentration determined by BCA assay. The samples were then diluted in Qiagen “wash buffer” containing 0.25% (v/v) CHAPS to a concentration of 0.1mg/ml. Aliquots of whole cell lysate and diluted whole cell lysate were kept for subsequent immunoblot. The phosphoenrichment columns were washed with 6ml “wash buffer”, before the diluted samples were loaded onto the columns. Following two further washes of the columns with 6ml “wash buffer”, the phosphoenriched fraction was eluted from the columns using Qiagen “elution buffer” containing 0.25% (v/v) CHAPS. Following concentration of the eluted fraction to a volume of 200-300ml using 9k molecular weight cut-off concentrator columns (Thermo Fisher) with centrifugation 13000 rpm 30 minutes, protein concentration was measured by BCA. For mass spectrometry (MS) experiments, phosphoenriched lysates were stored at −80°C. Otherwise, whole cell lysate, diluted whole cell lysate and phosphoenriched lysates were processed for SDS-PAGE with NuPage LDS Sample buffer (Life Technologies) and 100mM dithriothreitol (Sigma), followed by heating at 70°C for 5 minutes. Samples were then frozen at −80°C until immunoblotting was performed

### Immunoprecipitation

Samples were washed twice in ice cold HBS and then lysed in 1000ml lysis buffer for 30 minutes at 4°C with end over end mixing (Cell Signaling Lysis Buffer 20mM Tris-HCL pH7.5, 1mM Na_2_EDTA, 1mM EGTA, 1% Triton, 2.5mM sodium pyrophosphate, 1mM beta-glycerophosphate, 1mM Na_3_O_4_, 1mg/ml leupeptin supplemented with 1% (v/v) HALT protease inhibitor cocktail (Thermo Fisher) and 1% (v/v) Phosphatase inhibitor cocktail 2 and 3 (Sigma) and 1mM PMSF (Cell Signaling)). Lysates were clarified by centrifugation at 14000g for 15 minutes at 4°C, and the supernatant transferred to fresh Eppendorfs. Protein concentrations were calculated by BCA. 50ml of input lysate was heated with LDS/DTT and stored at −80°C for later immunoblot. Next, concentrations were adjusted to 1mg/ml and 7.5mg of protein was taken forward for IP for each condition. The appropriate antibody or isotype control antibody was added to the lysates followed by incubation with gentle end over end mixing at 4°C overnight.

The next morning, Protein G Dynabeads (Thermo Fisher) were washed once in lysis buffer, using a DynaMag2 magnet (Invitrogen) to separate the beads from solution, and 5µl of beads per 1µg of antibody was added to each sample. Samples were incubated with the beads with gentle end over end mixing for 2 hours at 4°C. Following this, the supernatant was removed using a magnet to separate the beads, with 50µl of the supernatant heated with LDS/DTT and stored at −80°C later immunoblot. The beads were washed 4 times in total with lysis buffer containing all protease and phosphatase inhibitors, with gentle end over end mixing for 5 minutes at 4°C for each wash. Elution of the beads was then performed by incubating the beads with pH 2.8 elution buffer (Pierce) with gentle end over end mixing for 30 minutes at 4°C. The eluate was collected and neutralised immediately with 1/10 volume of 1M Tris-HCL pH 9. Elution and neutralisation was performed a further two times to ensure complete elution. The eluate was stored at −80°C until used for downstream processing.

### Tandem Ubiquitin Binding Entities (TUBEs) Ubiquitin Immunoprecipitation

Typically 2.5×10^7^ THP-1 cells were used per condition. 30 minutes before the end of the experimental conditions, samples were incubated with 10µM MG132 (Sigma) for 30 minutes at 37°C before harvesting, washing once in ice cold PBS and lysing in 1ml Ub-IP lysis buffer (50mM Tris-HCL (pH 8.0), 150mM NaCl, 5mM EDTA, 1% NP-40, 0.5% Deoxycholate, 0.1% SDS, protease inhibitor cocktail (Roche), 1% (v/v) phosphatase inhibitor cocktail 3 (Sigma), 20µM MG132, 50µM PR619 and 100mM N-ethylmaleimide (Sigma). Cells were lysed for 30 minutes at 4°C with gentle end over end mixing. The lysate was clarified by centrifugation at 13000rpm for 20 minutes at 4°C, and the supernatant transferred to fresh Eppendorfs. Protein concentration was calculated by the Bradford assay and protein concentrations normalized to 2mg/ml. Typically 2mg of protein was taken forward for TUBEs IP. 50ul of input fraction was heated with LDS/DTT and stored at −80°C for later immunoblot. Samples were incubated with 40µl TUBE1 agarose beads (LifeSensors), or control agarose beads (LifeSensors) for 4 hours at 4°C with gentle end over end mixing. The beads were then centrifuged at 3000g for 3 minutes and washed with lysis buffer containing all inhibitors three times (each wash was performed for 5 minutes at 4°C with gentle end over end mixing), followed by two final washes with lysis buffer without SDS and Deoxycholate. Elution was then performed with 50ul of pH 2.8 elution buffer (Pierce) for 30 minutes with gentle end over end mixing at 4°C, followed by immediate neutralisation with 1/10 volume 1M Tris-HCL pH 9. Elution and neutralisation was performed three times in total. Samples prepared for later MS analysis were frozen at −80°C until further processing (100µl of 150µl total eluate). The remaining 50µl, and all eluate from other experiments were heated with 2xLDS/DTT at 70°C for 10 minutes and stored at −80°C until immunoblot.

### Liquid chromatography tandem mass spectrometry (LC-MS/MS)

100µl of lysate was adjusted to 175µl with ddH_2_O. All samples were then successively reduced and alkylated for 30 minutes with 5mM dithiothreitol and 20mM iodoacetamide, respectively. The proteins were then precipitated using chloroform-methanol precipitation and the pellet were solubilized in 6M urea, 0.1M Tris pH 7.8. The sample were diluted to 1M Urea. The digestion was performed overnight at 37°C by adding 500ng of trypsin. The peptides were desalted using a C18 cartridge (Waters). Briefly, the samples were conditioned with buffer A (1% (v/v) acetonitrile, 0.1% (v/v) trifluoroacetic acid (TFA) in water) prior to equilibration with buffer B (65% (v/v) acetonitrile, 0.1% (v/v) TFA in water). The acidified peptides were loaded onto the column, washed with buffer A and eluted with buffer B. The solution containing the peptides was dried with a speedvac and solubilised in 1% (v/v) acetonitrile, 0.1% (v/v) TFA in water for mass spectrometry analysis.

Peptides were analysed with nano ultra-high performance liquid chromatography tandem mass spectrometry (nano-UPLC-MS/MS) using a Dionex Ultimate 3000 nanoUPLC, coupled to an Orbitrap Fusion Lumos mass spectrometer (Thermo Scientific). MS analysis was performed essentially as described previously (84). In brief, the data were acquired with a resolution of 120,000 full-width half maximum at mass/charge 200 with EASY-IC using the reagent ion source (202 m/z) for internal calibration enabled, Top speed precursor ion selection, Dynamic Exclusion of 60 seconds and fragmentation performed in Collision Induced dissociation (CID) mode with Collision Energy of 35.

### Analysis of mass spectrometry data

#### Label-free quantitative analysis

The raw MS data were analysed using Progenesis QI (Waters) and searched using Mascot 2.5 (Matrix Science). The search settings were as follows: trypsin with 1 miscleavage allowed, oxidation (M) and Deamidation (N, Q) were set as variable modifications and carbamidomethylation (C) as fixed modification. The data was searched against human protein sequences using the UPR_homoSapiens_20141015 (85,889 sequences; 33,866,397 residues) allowing a peptide mass tolerance of 10 ppm and a fragment mass tolerance of 0.05 Da.

### Peaks search for phosphorylation

The raw data was analysed in PEAKS Studio 7.5 (Bioinformatics Solutions Inc). The settings were the following: The database used was the swissprot human database was used for the proteins identification. The enzyme used for the search was trypsin allowing a maximum of 2 miscleavages. Fixed modifications: Carbamidomethyl (C); Variable modifications: Deamidated (N), Deamidated (Q), Oxidation (M), Phospho (STY). 10 ppm mass tolerance were allowed for the precursor ions and 0.05 Da was allowed for the fragment ions.

### Mascot search for phosphorylation

The raw data was searched using Mascot with following settings. Enzyme: Trypsin; Fixed modifications: Carbamidomethyl (C); Variable modifications: Deamidated (N), Deamidated (Q), Oxidation (M), Phospho (ST), Phospho (Y); Peptide mass tolerance: ± 10 ppm (# 13C = 1); Fragment mass tolerance: ± 0.5 Da; Max missed cleavages: 1.

### ShRNA lentiviral transduction and siRNA transfection

Short hairpin RNA lentiviral particles were produced and transduced following the RNAi Consortium (TRC) protocols. ShRNA containing pLKO.1 vectors targeting NOD2 (SHCLND-NM_022162), ataxin-3 (SHCLND-NM_004993), TBK1 (SHCLND-NM_013254) or non-Target shRNA Control Plasmid DNA were all obtained from Sigma (MISSION shRNA Plasmid DNA). In brief, HEK293T packaging cells growing in 6 cm well plate were transfected with a mix of 1 µg packaging vector (psPAX2), 0.4 µg envelope vector (pMD2.G) and 1.6 µg hairpin-pLKO.1 vector (SHC016 control or gene specific shRNA. Fugene-6 (Promega) was used as transfection reagent. Cell culture medium containing lentiviral particles (LVP) was collected 48 h later and passed through a 0.45 µm filter (Sartorius). Virus preparations were then concentrated by centrifugation at 30,000 rpm for 90 min. Viral particles were added to cultured THP-1 cells in R10 (Roswell Park Memorial Institute medium (RPMI-1640) (Sigma) supplemented with 10% (v/v) heat-inactivated foetal calf serum (FCS) (Sigma), 2mM (1% v/v) L-glutamine (Sigma)) together with 8 µg/ml Polybrene (Sigma) toimprove transfection efficiency. Following incubation for 3 hrs at 37°C, the cells were harvested, washed, and resuspended at 1×106 cells/ml in R10 media with antibiotics including puromycin (as selective antibiotic). After 10 days of continuous selection with puromycin, knockdown efficiency was assessed by immunoblot. Transfection of human dendritic cells was performed by electroporation of SMARTpool ONTARGETplus human ataxin-3 (ATXN3) or non-targeting siRNAs (Dharmacon). Cells were resuspended in the solution provided with the kit (Invitrogen) followed by electroporation with Neon System kit (Invitrogen) using the following parameters: 1475 V, 20 ms, 2 pulses. After 48hrs, cells were harvested for use in experiments and to check knockdown by immunoblot.

### Adherent cell transfection

Human HEK293/NOD2 Cells were seeded 24 hours prior to transfection in media without antibiotics. Transfection mixes were made, comprising Fugene (Promega) at a ratio of 3:1 to amount of DNA plasmid to be transfected, in the appropriate volume of Opti-MEM (Gibco, Thermo Fisher) (10% of volume of media in wells to be transfected). The transfection mixes were incubated at room temperature for 20 minutes and then added dropwise to the wells to be transfected. Cells were either cultured for a further 24 or 48 hours before being used for downstream applications.

### RNA isolation

Typically 2-5 × 10^6^ cells per condition were harvested and washed once with cold PBS. Pellets were resuspended in 350µl RLT buffer (Qiagen) containing 1% (v/v) Mercapto-ethanol (Sigma) and stored at −80°C. Samples were thawed on ice and homogenized by adding to Qiashredder columns (Qiagen) and centrifuged 2 minutes 13000rpm. RNA isolation was then peformed using RNeasy kits (Qiagen) according to manufacturer’s instructions. The isolated RNA was eluted by added 25ml nuclease free water (Ambion) to the RNeasy column membrane for 5 minutes, followed by centrifugation into fresh Eppendorfs 8000g 1 minute. RNA concentration and purity were obtained using a Nanodrop 1000 spectrophotometer (Thermo Fisher) and samples were stored at −20°C until further analysis.

### Reverse Transcription

RNA was reverse transcribed using a high capacity RNA to cDNA kit (Applied Biosystems). 500ng to 2µg of RNA was normalized to the same concentration for each sample using nuclease free water (Ambion) in polypropylene PCR tubes (Starlab). Then an RT mix containing 2µl 10x RT buffer, 0.8µl 25x dNTP mix, 2µl RT random primers, 1µl multiscribe RT, 1µl RNase inhibitor (Applied Biosystems) and 3.2µl nuclease free water was added to each PCR tube (10µl total RT mix) (Starlab) for a total volume of RNA sample and RT mix of 20µl. This was reverse transcribed using a Thermo Cycler (Applied Biosystems) with the program: 25°C 10 minutes, 37°C 120 minutes, 85°C 5 minutes. The cDNA was stored at −20°C.

### Quantitative real time polymerase chain reaction (qPCR)

qPCR was performed using TaqMan chemistry (Applied Biosystems). cDNA was diluted 10 fold with nuclease free water. 4.5µl of diluted cDNA was added in triplicate for each sample to wells of a white 0.2ml 96 well PCR microplate (Starlab). 0.5ml of TaqMan FAM-MGB labelled primer (Applied Biosystems) and 5µl TaqMan Universal PCR Mastermix (Applied Biosystems) was added to each well, resulting in a 10ml total reaction mix. The plate was covered with a polyolefin optical film (Starlab) and centrifuged at 400g for 1 minute. qPCR was then performed using the Bio-Rad C1000 Thermal cycler CFX Realtime system (Bio-Rad) using the manufacturer’s recommended program: 50°C 2 minutes, 95°C 10 minutes, then 40 cycles of 95°C 15 seconds, 60°C 1 minute. Mean cycle threshold (Ct) number was calculated from the triplicate values. Relative gene expression was calculated in comparison to the housekeeping RPLP0 control. The difference in gene expression between conditions was calculated using the 2^−ΔΔCt^. This is derived from:

ΔCT=CT(target gene)-CT(control gene)

ΔΔCT=ΔCT(target condition)-CT(control condition)

### Flow cytometry

Typically 0.5×10^6^ THP-1 cells per condition were plated in 1ml of media in 12 well plates and differentiated for 16 hours with 25ng/ml PMA (Sigma). Following differentiation, the indicated treatments were applied. Cells were then harvested with gentle scraping and transferred to a FACS tube. The following staining protocols were then followed. Cytofluorometric evaluation was by the LSRII flow cytometer (BD Biosciences) with analysis of the data by FLOWJo.

### MitoSOX Red staining

The cells were pelleted by centrigugation and resuspended in room temperature HBSS (Thermo Fisher) to wash, then centrifuged. The cells were resuspended in 200µl MitoSOX red solution (final concentration 5µM MitoSOX red (Invitrogen) inHBSS) and incubated for 15 minutes in the 37°C cell culture incubator. 200µl of HBSS was added, and the tube centrifuged. The cells were resuspended in 250µl HBSS and cytofluorometric evaluation performed.

### Dihydrorhodamine 123 (DHR) staining

The cells were pelleted by centrifugation and resuspended in room temperature PBS to wash, then centrifuged. The cells were resuspended in 50µl DHR solution (final concentration 2.5µg/ml DHR 123 (Invitrogen) in PBS) and incubated for 30 minutes at 37°C in a water bath. As a positive control, 50µl of PMA (final concentration 100ng/ml (Sigma) in PBS) was added to control samples for the final 15 minutes. The cells were centrifuged and then washed once with PBS, before resuspension in 200µl FACS staining buffer. Cells were fixed with the addition of 200µl 1% PFA and cytofluorometric evaluation performed.

### Seahorse mitochondrial stress assay

The Seahorse XFe96 Extracellular Flux Analyser was used to measure mitochondrial respiration and glycolysis (Seahorse Bioscience). Seahorse 96 well assay plates (Seahorse Bioscience) were coated with Cell-Tak suspension (Corning). THP-1 cells were seeded at 1.5×10^5^ cells/well. Next, the optimal working concentrations of the compounds used for the mitochondrial stress test (oligomycin, FCCP and antimycin/rotenone) and the glycolysis stress test (oligomycin and 2-DG) were determined for THP-1 cells. The aim was to maximise the response to each compound with the lowest concentration possible. An XFe96 sensor cartridge (Seahorse) was hydrated overnight prior to Seahorse assays by adding 200ml of XF calibrant solution to each well and incubating in a CO2 free incubator at 37°C. The sensor cartridge (Seahorse) was loaded with the test drug compounds immediately prior to the assay and loaded on the Seahorse Analyser. 24 hours before the assay was run, cells were harvested, counted and resuspended at 1.88×10^6^ cells/ml. 2ml of cell suspension was plated in 6 well plates for each condition, with or without the appropriate ligand stimulation for the required duration. On the day of the assay, 1ml of cell suspension from each condition was harvested into 1.5ml Eppendorf tubes. For the mito stress test, cells were resuspended in mito stress test media with or without ligand(s) (XF base media (Seahorse Bioscience) supplemented with 1mM sodium pyruvate (Sigma), 5mM glucose (Life Technologies) and 2% FCS (Sigma) adjusted to pH 7.4 at 37°C and sterile filtered). 80µl of cell suspension (1.5×10^5^ cells) was then seeded in quadruplicate for each condition and the plate was centrifuged at 200g for 1 minute. Following 30 minutes in a CO_2_ free incubator at 37°C, 95ul of fresh mito stress test media was added to each well, and after a further 15 minutes in a CO_2_ free incubator the assay was run. Final concentrations of injected drugs were 1µM oligomycin, 1µM FCCP, 0.3µM rotenone and 0.3µM antimycin.

### STED immunofluorescence microscopy

Cells were plated in 8-well detachable tissue culture chambers on a PCA slide (Sarstedt) coated with 0.01% poly-l-lysine (Sigma) – cells were at a density of 1-2 × 10^5^ cell per well in 250ul of appropriate media. At the end of the experiment, cells were washed twice with PBS, fixed with 4% paraformaldehyde (Sigma) for 15 minutes and permeabilised with 0.5% (v/v) Triton X-100 (Sigma). Cells were blocked overnight at 4°C with 150µl blocking solution per well (5% (v/v) human serum (Sigma), 5% (v/v) goat serum (Sigma), 5% (v/v) FCS (Sigma). The following day, cells were incubated with primary antibody diluted in 140µl blocking solution at a pre-optimised or manufacturer recommended concentration for 1 hour at room temperature. Following three washes with PBS (250µl per well, 5 minutes gentle shaking), cells were incubated with the species appropriate fluorescently labelled secondary antibody for 1 hour at room temperature. Cells were washed three further times with PBS, and the detached slide was then mounted with Vectashield mounting media containing DAPI (Vector Laboratories). Alternatively, cells were incubated with 200µl PBS containing 1:100 DAPI (Thermo Fisher) for 15 minutes at room temperature and the detached slide was mounted with Vectashield mounting media without DAPI (Vector Laboratories). A Leica SP8 STED system was used for imaging. ImageJ was used for image processing and analysis.

### Bacterial killing assay

1×10^6^ THP-1 cells were seeded per condition in 1ml R10 without antibiotics in a 24 well plate, and differentiated overnight with 25ng/ml PMA. After 16 hours, the media was changed for 500µl fresh R10 media without antibiotics. Two hours later, *Salmonella Typhimurium* LT2 was added at a multiplicity of infection (MOI) of 20:1. Thirty minutes post infection, wells were washed twice with PBS and 500ml fresh R10 supplemented with Gentamicin 100µg/ml was added. After a further thirty minutes, wells were washed once with PBS and 500µl fresh R10 supplemented with Gentamicin 30µg/ml was added. At the end of the designated post infection period, the medium was removed (and stored at −80°C until later analysis by ELISA) and 500µl of PBS with 1% (v/v) saponin was added to the wells, followed by incubation for 5 minutes at 37°C. 500µl of PBS was added and serial dilutions plated on LB/agar plates, incubated overnight at 37°C, and colonies then counted. Alternatively, when cell viability post infection was assessed, cells were detached by incubating with 500µl of trypsin per well for 5 minutes at 37°C, then viability assessed by trypan blue staining (Invitrogen) and counting of live/dead cells.

### Statistical analysis

Prism (GraphPad) was used to determine the statistical significance. When making multiple comparisons on a data set, analysis was by one-way ANOVA with post-hoc Bonferroni analysis. For experiments with two sample groups (one condition, one control) and a single comparison, analysis was by paired, two-tailed Student’s t-test. Error bars represent Standard Error of the Mean (SEM).

## Supplementary information

**Supplementary Figure 1 (S1).**
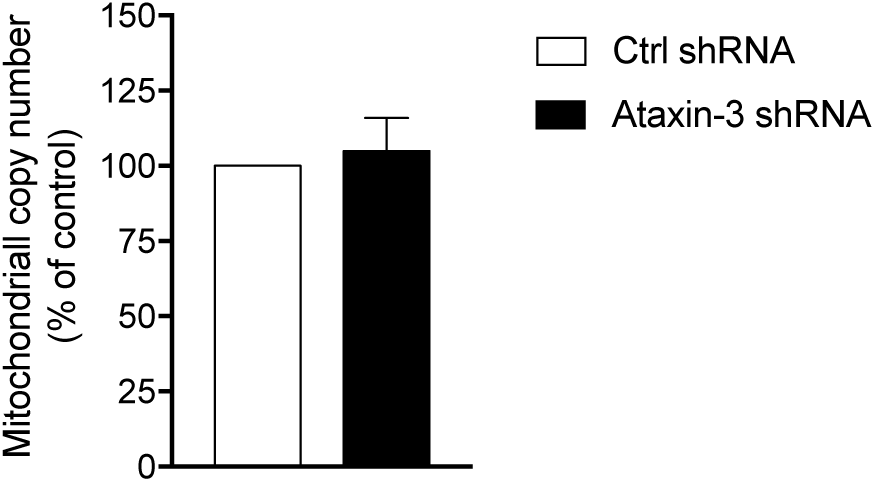
SYBR Green based qPCR of a mitochondrial DNA fragment (mitochondrially encoded tRNA leucine 1) was performed, and the relative mtDNA copy number calculated by normalising to simultaneous qPCR of a nuclear DNA fragment (b2 microglobulin). n=2.

**Supplementary Figure 2 (S2).**
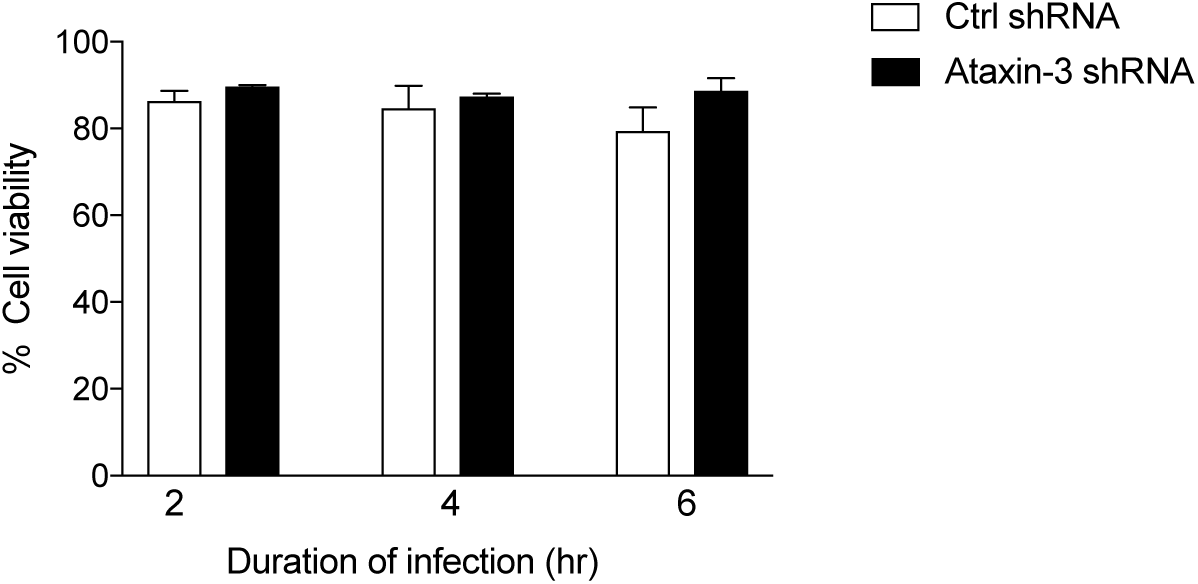
THP-1 cell viability in ataxin-3 and control shRNA THP-1 cells infected for 1-6hr with *Salmonella enterica* serovar Typhimurium. n=3.

**Supplementary Table 1.** Proteins found to be differentially phosphorylated in phosphoproteomic analysis of human moDCs following MDP+PAM_3_CSK_4_ stimulation

| Protein (Gene) | Log2 fold change |
| --- | --- |
| Ubiquitin-like modifier-activating enzyme 6 (UBA6) | 5.36 |
| Coagulation factor X (F10) | 4.70 |
| GA-binding protein alpha chain (GABPA) | 4.57 |
| DnaJ homolog subfamily B member 1 (DNAJB1) | 4.02 |
| Ubiquitin thioesterase otulin (OTULIN) | 3.88 |
| THAP domain-containing protein 11 | 3.36 |
| ADP-ribosylation factor-binding protein GGA1 (GGA1) | 3.12 |
| TNF receptor-associated factor 6 (TRAF6) | 3.03 |
| BRCA1-associated ATM activator 1 (BRAT1) | 2.95 |
| Trifunctional purine biosynthetic protein adenosine-3 (GART) | 2.91 |
| Dual specificity mitogen-activated protein kinase kinase 6 (MAP2K6) | 2.88 |
| Ataxin-3 (ATXN3) | 2.86 |
| Myeloma-overexpressed gene 2 protein (MYEOV2) | 2.74 |
| Glutamine-dependent NAD(+) synthetase (NADSYN1) | 2.72 |
| Zinc finger MIZ domain-containing protein 1 (ZMIZ1) | 2.71 |
| Bromodomain-containing protein 4 (BRD4) | 2.68 |
| Histone deacetylase 6 (HDAC6) | 2.63 |
| Leucine-rich repeat-containing protein 41 (LRRC41) | 2.58 |
| Vacuolar-sorting protein SNF8 (SNF8) | 2.48 |
| Vacuolar protein sorting-associated protein VTA1 homolog (VTA1) | 2.37 |
| Beta-adrenergic receptor kinase 1 (ADRBK1) | 2.34 |
| Target of Myb protein 1 (TOM1) | 2.34 |
| Endophilin-B1 (SH3GLB1) | 2.27 |
| Translocon-associated protein subunit beta (SSR2) | 2.21 |
| Protein S100-A6 (S100A6) | 2.19 |
| AN1-type zinc finger protein 5 (ZFAND5) | 2.18 |
| Short coiled-coil protein (SCOC) | 2.18 |
| Signal transducer and activator of transcription 6 (STAT6) | 2.16 |
| Signal transducer and activator of transcription 1-alpha/beta (STAT1) | 2.13 |
| 26S proteasome non-ATPase regulatory subunit 4 (PSMD4) | 2.10 |
| Coagulation factor IX (F9) | 2.07 |
| ADP-ribosylation factor-binding protein GGA2 (GGA2) | 2.06 |
| Syntenin-1 (SDCBP) | 2.04 |
| Dual specificity mitogen-activated protein kinase kinase 3 (MAP2K3) | 2.03 |
| Vitronectin (VTN) | 2.01 |
| Protein NEDD1 (NEDD1) | 2.00 |
| Mitogen-activated protein kinase 14 (MAPK14) | 1.97 |
| 2'-deoxynucleoside 5'-phosphate N-hydrolase 1 (DNPH1) | 1.95 |
| Proteasome inhibitor PI31 subunit (PSMF1) | 1.93 |
| CTD small phosphatase-like protein 2 (CTDSPL2) | 1.90 |
| Anamorsin (CIAPIN1) | 1.90 |
| Proteasomal ubiquitin receptor ADRM1 (ADRM1) | 1.89 |
| Tristetraprolin (ZFP36) | 1.89 |
| NLR family CARD domain-containing protein 4 (NLRC4) | 1.87 |
| Heat shock protein beta-1 (HSPB1) | 1.87 |
| Coronin-7 (CORO7) | 1.85 |
| Hyaluronan-binding protein 2 (HABP20) | 1.84 |
| Phosphatidylinositol-binding clathrin assembly protein (PICALM) | 1.79 |
| Receptor-interacting serine/threonine-protein kinase 1 (RIPK1) | 1.78 |
| Nipped-B-like protein (NIPBL) | 1.74 |
| Exportin-1 (XPO1) | 1.73 |
| Host cell factor 1 (HCFC1) | 1.70 |
| AN1-type zinc finger protein 6 (ZFAND6) | 1.69 |
| Golgi reassembly-stacking protein 2 (GORASP2) | 1.69 |
| Protein FAM161B (FAM161B) | 1.65 |
| Tensin-like C1 domain-containing phosphatase (TENC1) | 1.63 |
| Stathmin (STMN1) | 1.62 |
| ATP-dependent RNA helicase DHX29 (DHX29) | 1.61 |
| Vacuolar protein-sorting-associated protein 25 (VP25) | 1.61 |
| Dual specificity mitogen-activated protein kinase kinase 4 (MAP2K4) | 1.59 |
| TRAF family member-associated NF-kappa-B activator (TANK) | 1.59 |
| Ubiquitin carboxyl-terminal hydrolase 25 (USP25) | 1.56 |
| Ankyrin repeat domain-containing protein 13A (ANKRD13A) | 1.55 |
| Retinoic acid receptor RXR-alpha (RXRA) | 1.54 |
| Toll-interacting protein (TOLLIP) | 1.53 |
| Spartin (SPG20) | 1.53 |
| SEC23-interacting protein (SEC23IP) | 1.53 |
| Probable helicase senataxin (SETX) | 1.53 |
| Sorting nexin-12 (SNX12) | 1.52 |
| CREB-binding protein (CREBBP) | 1.50 |

**Supplementary Table 2.**
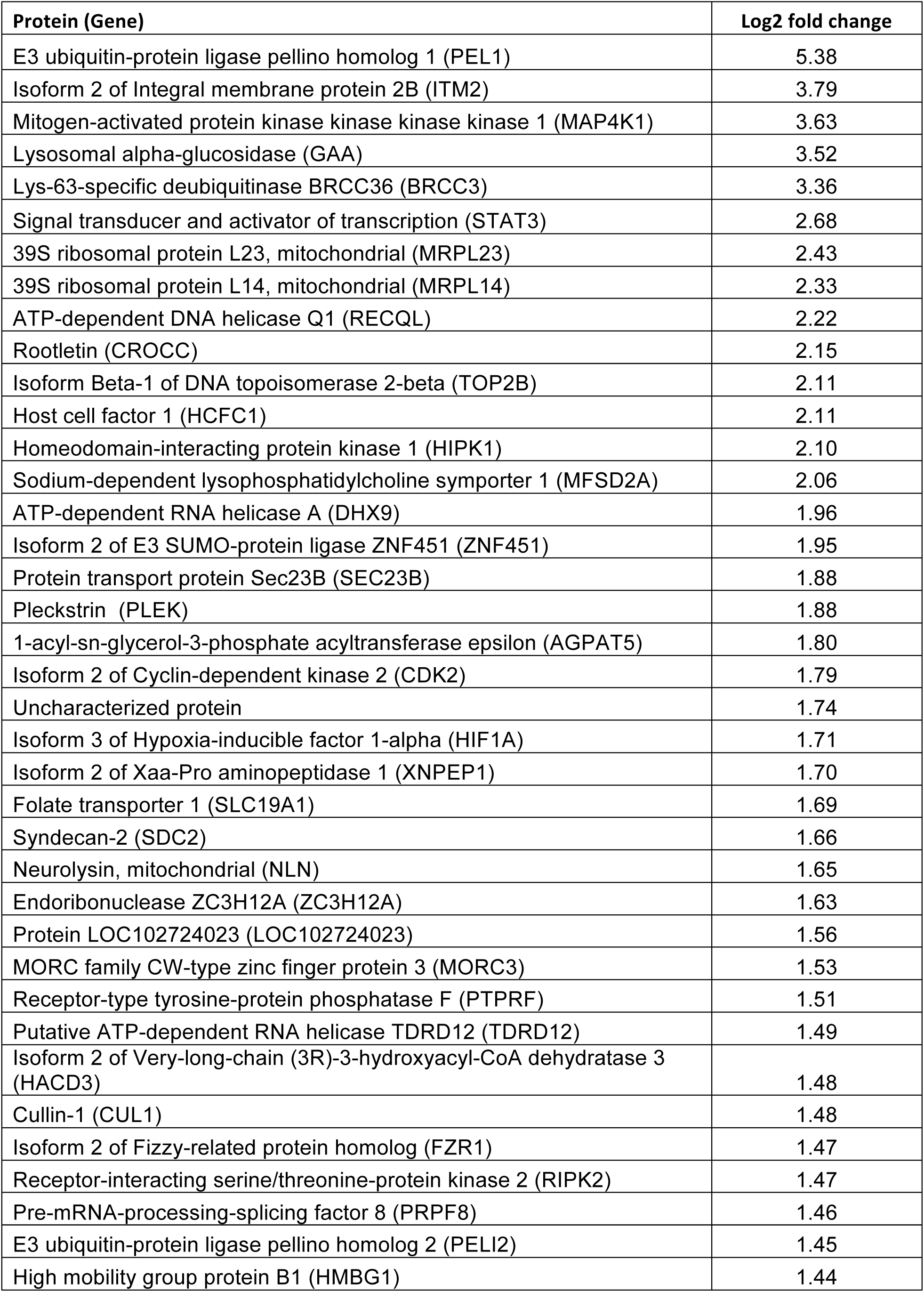
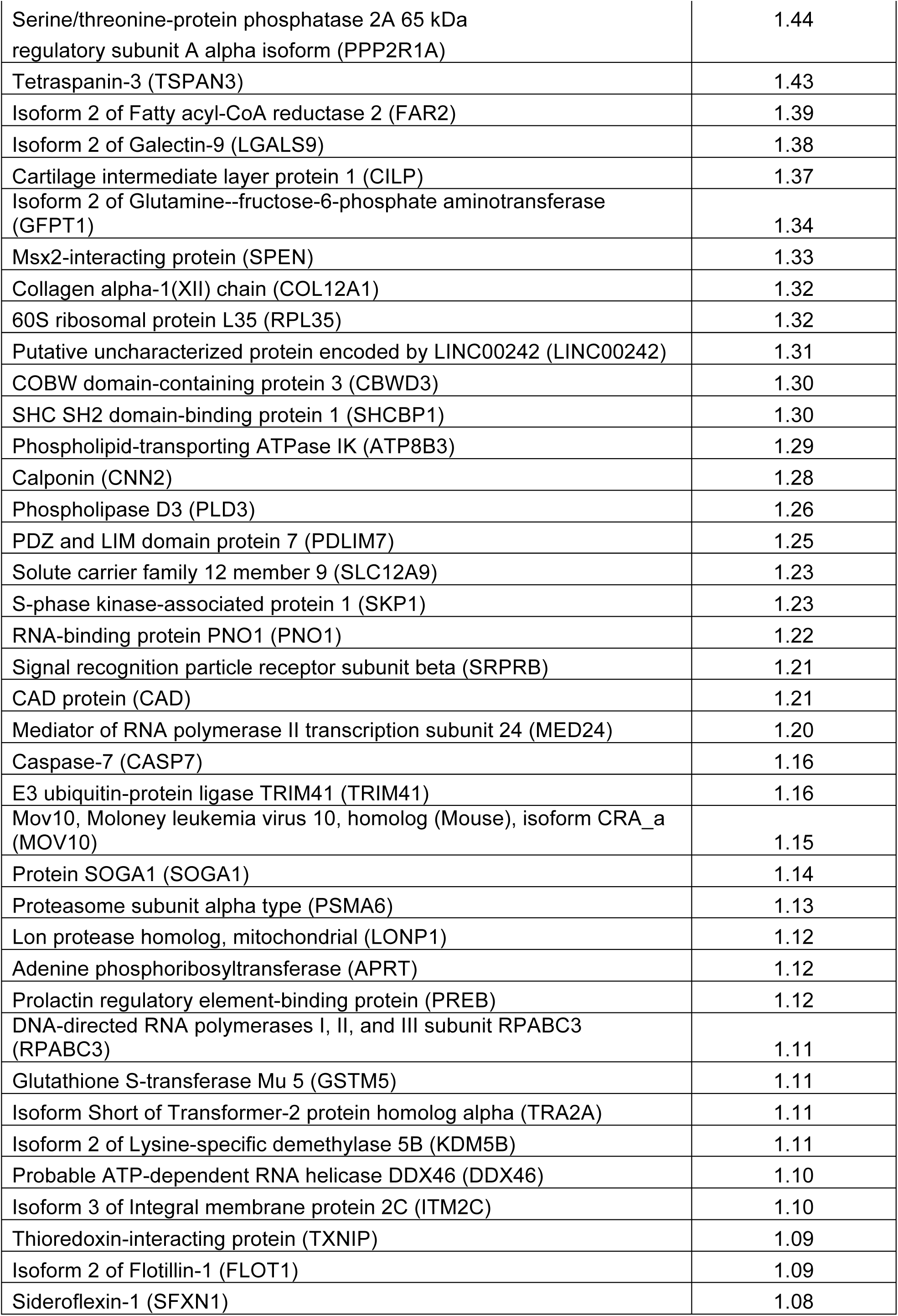
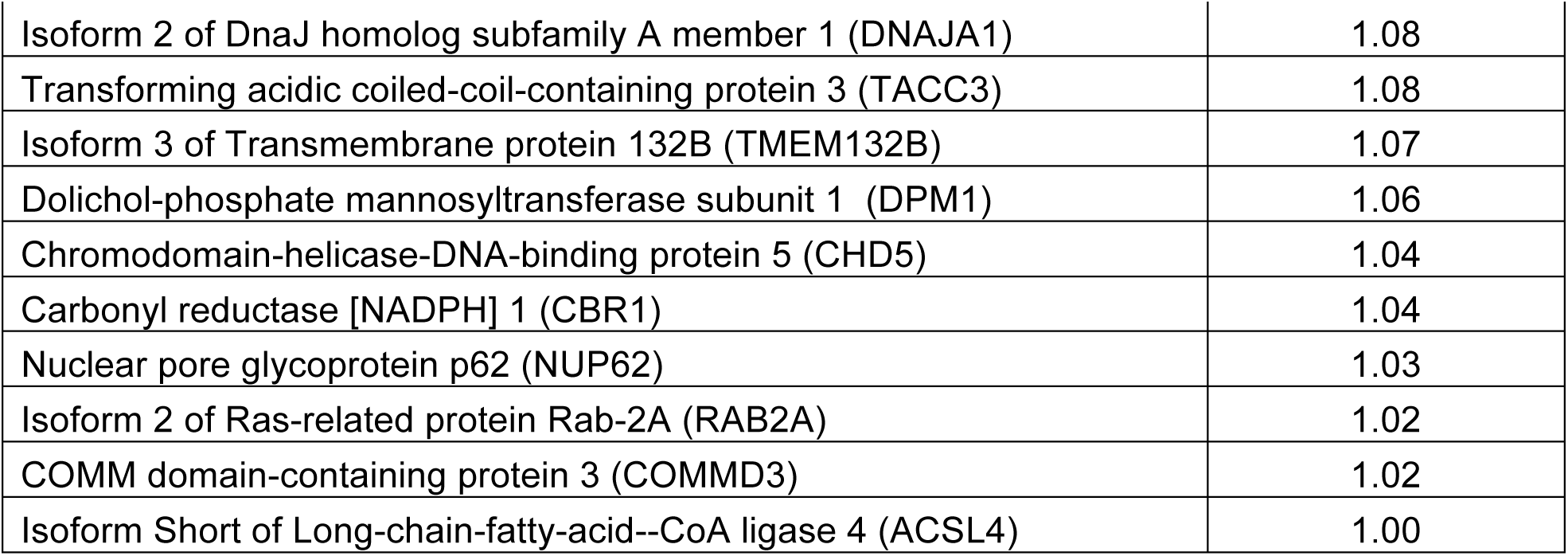
Proteins found to be differentially ubiquitinated in TUBES analysis comparing control and ataxin-3 depleted THP-1 cells

**Supplementary Table 3.**
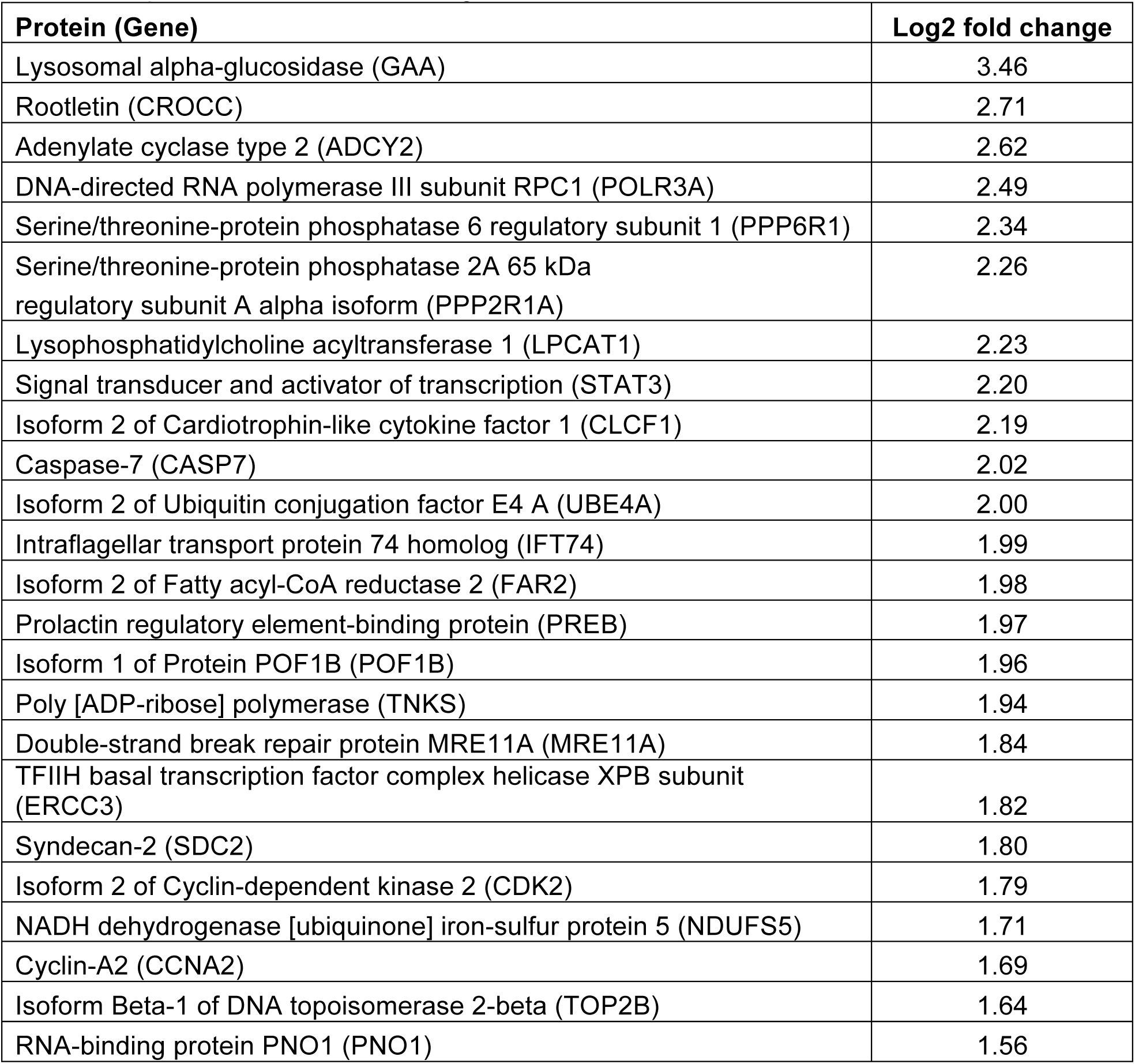
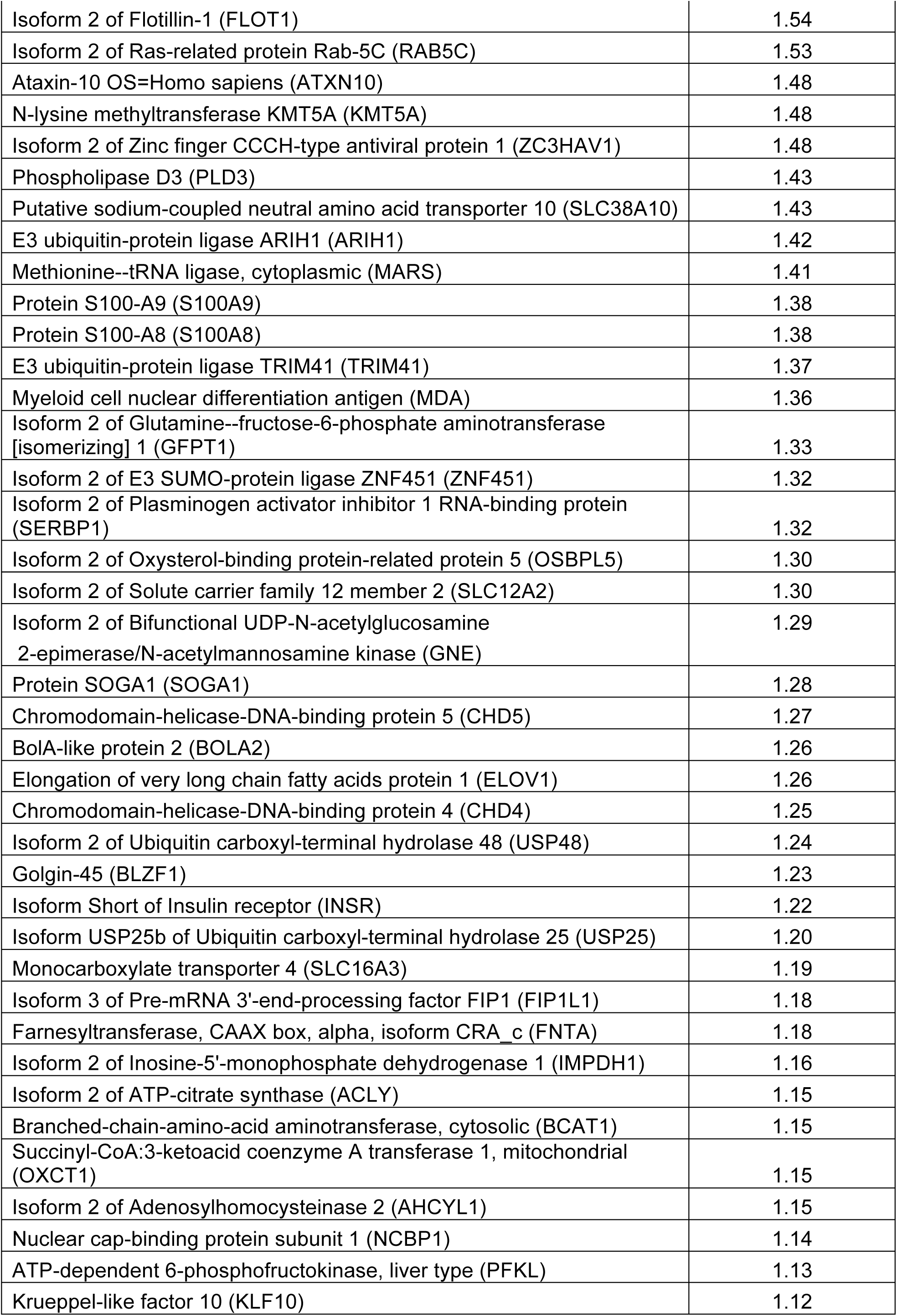
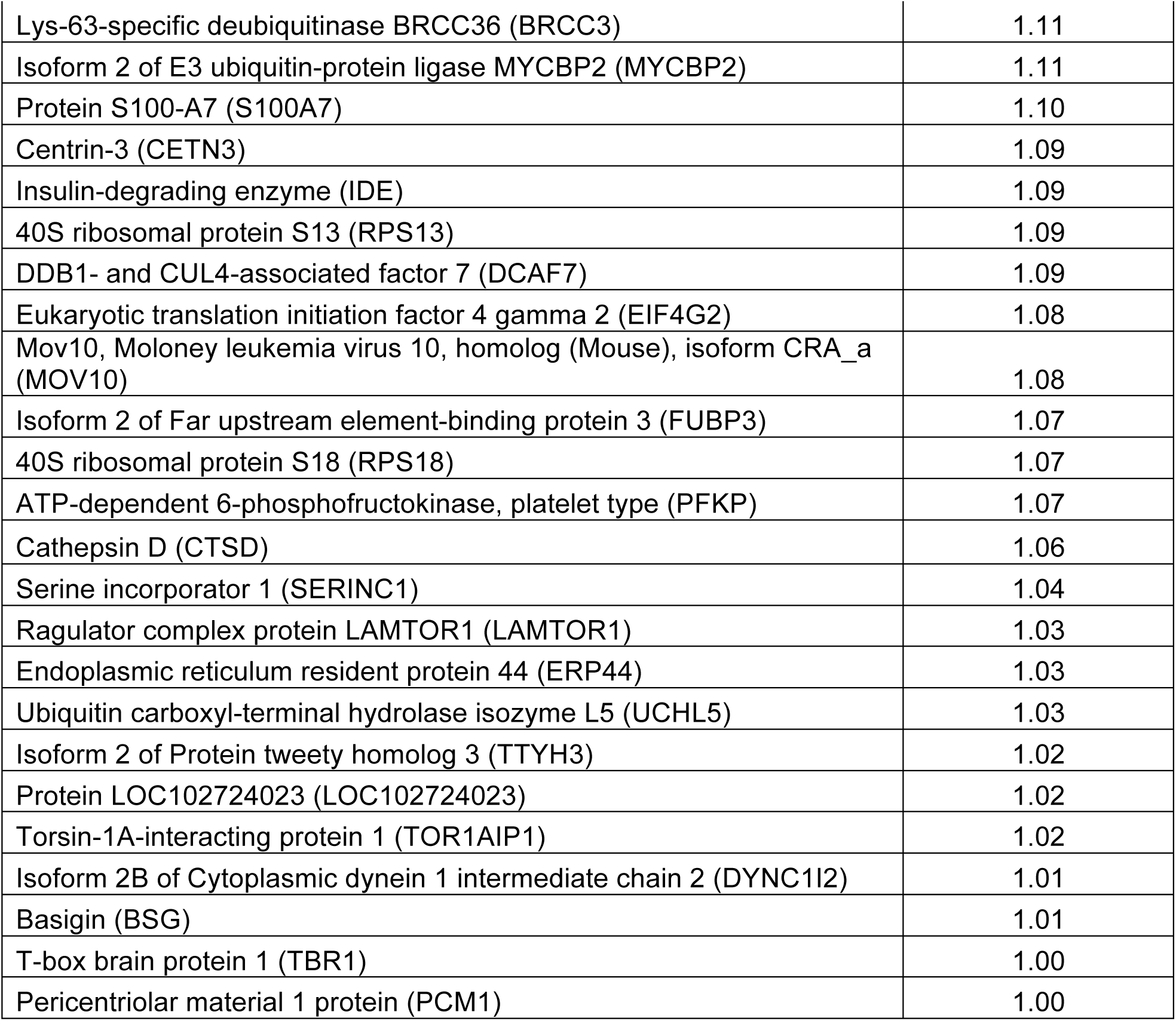
Proteins found to be differentially ubiquitinated in TUBES analysis comparing control and ataxin-3 depleted THP-1 cells following MDP+PAM_3_CSK_4_ stimulation

